# Environmental Noise and Lag Select Distinct Phenotypic Bet Hedging in Microbial Metabolic Allocation

**DOI:** 10.64898/2026.11.27.769635

**Authors:** Emanuele Pigani, Roaa M. Y. Omer, Onofrio Mazzarisi, Giuseppe Giuliani, Jacopo Grilli

## Abstract

Genetically identical microbial cells often differ in their growth rate and metabolic state, and in how long they take to resume growth after a shift of conditions. Preparing for uncertain future conditions is costly: cells that allocate part of their proteome to a future nutrient grow more slowly on the present one. Bet hedging offers a complementary strategy: a genotype whose cells occupy different metabolic states makes it likely that some of them are suited to whatever condition follows and resume growth quickly. Here, we investigate when phenotypic variability in metabolic allocation can outperform the evolutionary stable homogeneous strategy, in which all cells share the same allocation. We consider serial-dilution cultures, where each cycle ends when the nutrient is exhausted, and fluctuations in dilution or nutrient supply change the fold increase in biomass required in each cycle, i.e., the growth factor. We identify two distinct mechanisms by which selection can favor phenotypic heterogeneity. First, noise in the growth factor leaves the optimal mean allocation unchanged but can destabilize the homogeneous strategy: cycles that demand much growth favor fast growers, whereas cycles that demand little growth favor well-prepared cells. Variability in allocation is therefore favored above a critical noise level, set by the mean logarithm of the growth factor and by how steeply the lag increases as the allocation to the incoming nutrient decreases. Second, with multiple possible future nutrients, different cells can allocate their proteome to different future nutrients. When the lag is sufficiently long relative to the growth time, an all-in strategy, in which cells specialize in individual future nutrients, is favored independently of noise in the growth factor. Invasion analysis of narrow phenotypic distributions shows that these two mechanisms define four evolutionary regimes, confirmed by competition and evolution simulations. Our results show how environmental uncertainty can make cell-to-cell variability selectively advantageous, so that evolution acts on the distribution of phenotypes within a genotype, not only on its mean.

## I. INTRODUCTION

Microbial populations encounter environments that change in nutrient composition and, more generally, in physico-chemical conditions. These changes arise as resources are released, transported, and consumed, and as cells move between habitats. In the ocean, nutrient limitation constrains microbial activity, and cells experience a nutrient supply that fluctuates in time, alternating between abundance and scarcity, as they encounter patchy, transient sources such as phytoplankton [1–3]. Laboratory serial-transfer experiments impose a similar feast–famine structure, in which each transfer supplies fresh nutrients that the population then exhausts. In both settings, the conditions experienced by a lineage differ from those of a fixed growth medium, and adaptation must be assessed across the sequence of environments encountered over successive generations [4]. Because growth is multiplicative, long-term success depends on the growth accumulated over the whole sequence of environments, so the strategy that grows fastest in one environment need not be the most successful overall [5–7]. For microbes facing nutrient changes, this raises the physiological question of how investment in current growth should be balanced against the capacity to grow after the environment changes.

Environmental correlations can partly solve this problem: when some conditions reliably follow others, cells can prepare for what is likely to come next [8, 9]. When the future environment cannot be predicted reliably, a clonal population can instead spread its cells over phenotypes suited to different environments [10, 11]. Such diversification is bet hedging when it lowers the average growth of the genotype but increases its long-term growth across sequences of environments [12, 13]. The benefit accrues to the genotype as a whole, not to individual cells. In engineered yeast, the relative success of populations with different switching rates depends on how frequently environments change [14], and stochastic switching has evolved in laboratory evolution experiments with bacteria [15]. Preparing for likely futures and diversifying phenotypes are, therefore, two complementary, not mutually exclusive, strategies to address uncertainty about the environment. In this work, we introduce a framework for studying how these strategies coevolve.

The way cells allocate their limited proteome determines the costs and benefits of these strategies. The expression of proteins that do not contribute to current growth comes at a well-defined cost, quantified by bacterial growth laws: cells can increase their preparedness for future environmental conditions, i.e., the proteome already allocated to them, at the expense of their current growth rate [16]. This trade-off is further supported by nutrient-shift experiments and allocation models, which reveal that faster growth before a shift comes at the cost of a longer lag afterward [17–20].

Metabolic preparedness is therefore a quantitative trait on which selection can act [9, 19, 21]. Previous studies have typically assumed a homogeneous allocation strategy within a population, establishing a one-to-one corrspondence between genotype and phenotype, with each genotype encoding a specific balance between growth and lag. Here, we relax this assumption by allowing a single genotype to generate phenotypic heterogeneity, giving rise to a distribution of allocation strategies rather than a single, uniform allocation. Single-cell experiments provide evidence for such phenotypic heterogeneity, showing that differences in metabolic state before a nutrient shift can lead to differences in subsequent growth responses [22–24]. In *E. coli*, phenotypic heterogeneity can also emerge in response to the shift itself, giving rise to distinct growing and nongrowing subpopulations [25].

Proteome-allocation theory provides a natural starting point to ask when a distribution of allocation strategies outperforms the best single allocation. Previous work linked allocation to growth and lag in a resource-depletion model, showing how selection among homogeneous strategies balances these traits and sets how much proteome is allocated to future nutrients according to the probabilities of environmental transitions [9]. Related theory has identified conditions favoring mixtures of growing and arrested phenotypes along growth–lag trade-offs when the durations of environmental periods are specified externally [26]. However, in a serial dilution with resource depletion, each cycle lasts until the supplied nutrient is exhausted. The required expansion of biomass, which we call the growth factor, i.e., the ratio between the biomass at depletion and at the start of the cycle, is therefore set by the nutrient supply and by the biomass left by the previous dilution. Even when nutrients alternate deterministically, variable dilution or supply makes the growth factor fluctuate from cycle to cycle. A large growth factor favors fast growth, a small one favors a short lag. Whether such fluctuations can favor phenotypic diversification over the best homogeneous allocation has not been established.

Here, we extend the allocation model so that each geno-type expresses a distribution of allocation phenotypes. In our setting, the allocation phenotype itself is not heritable, whereas its distribution is, because it is genetically encoded. We ask when a genotype that generates heterogeneous phenotypes can outcompete the best homogeneous allocation.

## II. RESULTS

We model bacterial growth under serial dilution in fluctuating nutrient environments using a coarse-grained description of proteome allocation. The proteome is partitioned into housekeeping, ribosomal, and metabolic sectors, with mass fractions *φ*_*H*_, *φ*_*R*_, and *φ*_*P*_, respectively.

The housekeeping sector is fixed and includes the inactive ribosomes [9], so that *φ*_*R*_ counts only active ribosomes and *φ*_*H*_ + *φ*_*R*_ + *φ*_*P*_ = 1.

Metabolic allocation depends on the currently available nutrient: a fraction *f* of the metabolic sector is devoted to supporting growth in the current environment, while the remaining fraction 1 *− f* is preallocated among the proteins required under possible future nutrient conditions. Thus, *f* controls the trade-off between immediate growth and preparedness for subsequent environmental changes.

### A. Environmental noise makes bet hedging favorable over homogeneous allocation in two-resource environments

For simplicity, we begin with two nutrient conditions, *A* and *B*. The environment switches between these two conditions over successive serial dilution cycles (see Figure 1). The allocation to the currently available nutrient of a phenotypically homogeneous population, in which all cells express the same allocation strategy *f*, determines its exponential growth rate *λ*. The preparedness ratio *r* = (1 *− f* )*/f*, the allocation to the alternative nutrient relative to that to the current one, instead determines the lag time *T* incurred after a nutrient shift (see Figure 1A, more details in Material and Methods). Increasing *f* therefore accelerates growth in the current environment, but prolongs the lag following an environmental change. Together, growth and lag determine the time needed to complete the biomass expansion required in each dilution cycle: lag delays the onset of growth, whereas the exponential growth rate controls the subsequent population expansion. In this homogeneous limit, evolutionary dynamics select an optimal allocation *f*_opt_ that minimizes this resource-depletion time, thus balancing growth and lag [9].

**FIG 1.**
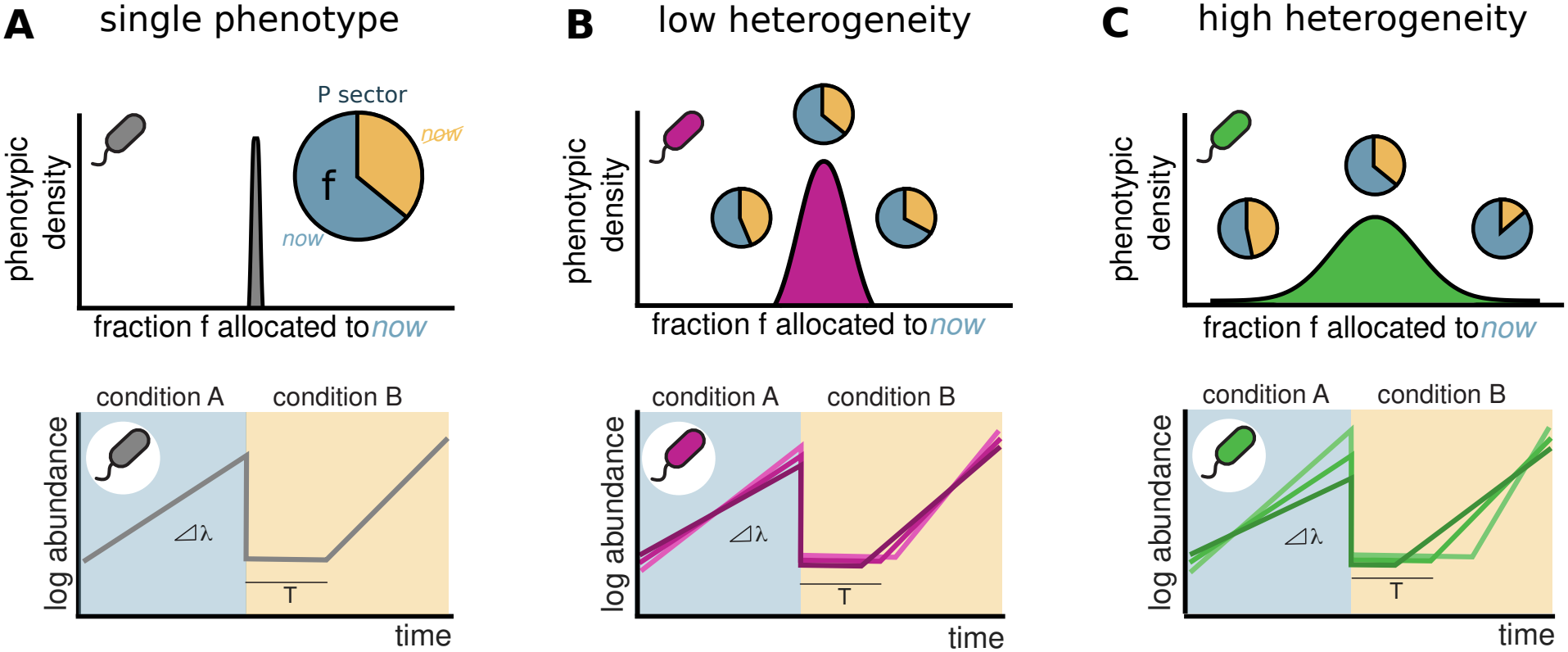
Metabolic bet hedging differs from homogeneous allocation to future resources. (A) A homogeneous genotype expresses one allocation phenotype: all cells strike the same balance between current growth and future preparedness. (B, C) A heterogeneous genotype instead distributes cells across allocation phenotypes around a mean state. The level of heterogeneity is a heritable trait that can vary (low in panel B and high in panel C) across genotypes. Predictable environments favor the best homogeneous allocation, whereas sufficiently noisy environments can favor a heterogeneous genotype because different cells are close to the phenotype favored in different cycles.

To account for phenotypic variability within clonal bacterial populations, we allow genetically identical cells to express different metabolic allocation strategies (see Figure 1B). The allocation values of identical cells vary around the genotypic mean *f*, with variability *σ*_*f*_ . When this variability is small, phenotype-specific growth rates and lag times remain close to those of the homogeneous population, and the population-level dynamics are nearly unchanged. Cells with larger *f* grow faster in the current environment but experience longer lags after a nutrient shift, whereas cells with smaller *f* exhibit the opposite behavior. As *σ*_*f*_ increases, these physiological differences become more pronounced, generating distinct growth-oriented and future-prepared phenotypes within the same genotype (see Figure 1C).

Relative to the optimal homogeneous allocation, phenotypic diversification carries a cost under a fixed growth factor because some cells express suboptimal allocation strategies. It is not immediately clear whether fluctuations in the growth factor can compensate for this cost and favor a nonzero *σ*_*f*_ . We therefore determine the conditions under which phenotypic diversification constitutes an advantageous bet-hedging strategy.

We first characterize the growth factor experienced in each serial dilution cycle. Let *N*^*d*^ denote the population biomass at the beginning of cycle *d*, and let *c*^*d*^ denote the supplied resource concentration, expressed in biomass equivalents. Without loss of generality, we assume yield equal to 1, so that complete resource depletion requires the population to expand from *N*^*d*^ to *N*^*d*^ + *c*^*d*^. After depletion, the population is diluted by a factor *D*^*d*^, such that the biomass at the beginning of the next cycle is *N*^*d*+1^ = (*N*^*d*^ + *c*^*d*^)*/D*^*d*^. We define the logarithmic growth factor *δ*^*d*^ of cycle *d* as the logarithm of the cycle growth factor, that is, of the increase in relative biomass before dilution, namely

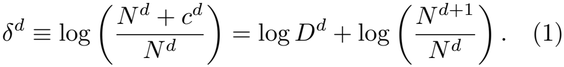

Thus, *δ*^*d*^ = log *D*^*d*^ in the deterministic stationary state, whereas the two quantities generally differ from cycle to cycle when dilution or resource supply fluctuates.

For a homogeneous phenotype with allocation *f*, the time required to complete this expansion is

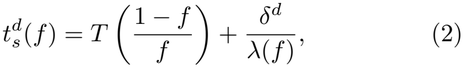

and is minimized by *f*_opt_ [9]. The first term is the lag *T* incurred after the nutrient shift, which is a function of the preparedness ratio *r* for the alternative nutrient, while the second is the time required for exponential growth.

Both the dilution factor *D*^*d*^ and the resource supply *c*^*d*^ may vary across serial dilution cycles: fluctuations in resource supply directly change the biomass expansion required for resource depletion, whereas fluctuations in dilution alter the biomass inherited by the following cycle. We write the cycle growth factor as 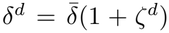 where 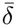 is its stationary mean and *ζ*^*d*^ encodes fluctuations, with zero mean at stationarity. We quantify the magnitude of growth-factor fluctuations by *σ*_*ζ*_ = sd_stat_(*ζ*^*d*^). At stationarity, the mean log growth factor equals the mean logarithmic dilution,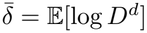.

For the sake of analytical simplicity, we first isolate dilution-driven variability of the growth factor by considering a fixed resource supply, *c*^*d*^ = *c*_0_, while allowing the dilution factor *D*^*d*^ to fluctuate across cycles, with dilution noise 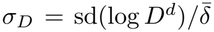. The two nutrient conditions continue to alternate, so that every cycle begins with a nutrient shift and a different dilution factor.

To begin with, we consider a resident genotype, in which all cells express the optimal homogeneous allocation *f*_res_ = *f*_opt_. This homogeneous resident is challenged by a mutant genotype that has the same mean allocation *f*_mut_ = *⟨f*_opt_*⟩* but distributes its cells around it with variability 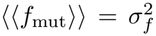. In other words, the mutant has no advantage in mean allocation: any difference in fitness can only come from the diversity of phenotypes it expresses.

We first simulate pairwise competitions between these two genotypes, starting from equal biomass, under increasing levels of dilution noise (see Figure 2A). When dilution noise is low, any level of phenotypic heterogeneity causes biomass losses in the mutant population and eventually leads to its exclusion, even when its mean allocation is optimal: in a nearly constant environment, spreading cells away from the optimum only carries a cost. When the dilution noise is sufficiently high, the outcome depends on how broad the mutant distribution is. A mutant with low to moderate variability can take over the population and does so in most stochastic histories. Instead, a mutant with excessive variability is excluded because too many of its cells express allocations that perform poorly in every cycle. Environmental noise can therefore favor phenotypic heterogeneity, but only above a certain level and only within a limited range of heterogeneity.

**FIG 2.**
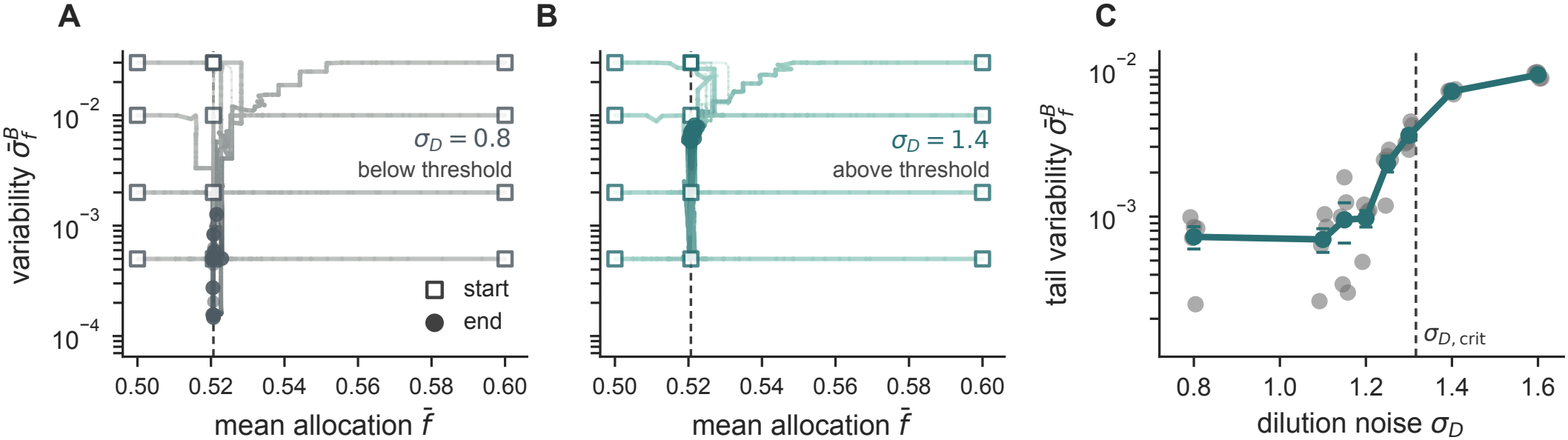
Environmental noise favors a heterogeneous genotype over a homogeneous one, up to a maximum variability. (A) Representative fixed-strategy competitions between a homogeneous resident at *f*_opt_ and heterogeneous mutants with the same mean allocation. The three lines of different colors correspond to different levels of phenotypic variability *σ*_*f*_ and of dilution noise *σ*_*D*_ . The colors and the markers next to the labels match the three points in panel B, and shaded regions show quantiles across stochastic histories. (B) Mutant win probability in the (*σ*_*D*_, *σ*_*f*_ ) plane. The dashed vertical line marks the local theoretical threshold for mean log growth factor 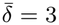 after converting from *σ*_*ζ*_ to *σ*_*D*_, the white curve marks the empirical finite-variability transition, and the colored markers mark the three competitions shown in (A). (C) Invasion of heterogeneous residents by rare heterogeneous mutants at *σ*_*D*_ = 1.5, both with mean allocation *f*_opt_. Color shows the invasion fitness *s* (see Eq. (34)), estimated as the change in mutant-to-resident log-biomass ratio per serial dilution cycle and averaged over stochastic histories, on a symmetric logarithmic scale. White curves mark *s* = 0: one branch follows the diagonal (dashed), where mutant and resident coincide and are neutral by construction; the other branch separates mutants that invade from those that are excluded. Their crossing marks the singular variability 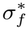 narrower residents are invaded by broader mutants, broader residents by narrower ones, and a resident at 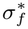 is invaded by neither.

We examine the probability that the mutant outcompetes the resident as a function of the dilution noise and the phenotypic variability (see Figure 2B). Below a critical dilution noise *σ*_*D*,crit_, the mutant loses at all levels of variability. Above threshold, heterogeneous mutants win for all levels of variability up to an upper limit 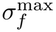, which increases with the noise (white curve). Narrow distributions also invade, but more slowly: they are nearly indistinguishable from the resident and need many cycles to take over, so that the transition is blurred at the lowest variabilities, where competitions are not concluded within the simulated time. Broad distributions are instead excluded because the cost of placing cells far from the optimum eventually dominates any benefit from fluctuations.

When the resident is itself heterogeneous (see Figure 2C), at high noise a rare mutant with larger variability invades a narrower resident, and a rare mutant with smaller variability invades a broader one, so selection converges toward intermediate variability rather than ever-broader distributions.

To determine *σ*_*D*,crit_, we perform an invasion analysis in which the mutant is initially rare, so that the resident alone sets the biomass dynamics and hence the duration of each cycle (see Material and Methods). The mutant invades if its invasion fitness, defined as the long-run mean change per cycle of the mutant-to-resident log-abundance ratio, is positive. Because the growth rate in Eq. (2) is a monotone function of *f*, it is convenient to use *λ* itself as the phenotypic coordinate and to write the lag as *T* (*λ*). The time needed to complete a cycle with log growth factor 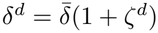 is then

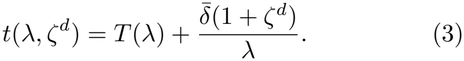

Since E[*ζ*^*d*^] = 0, the optimal homogeneous allocation minimizes the mean completion time and does not depend on the magnitude of growth-factor fluctuations. Mutant subpopulations have growth rates *λ*_opt_ + *ϵ*, where *λ*_opt_ = *λ*(*f*_opt_), with *⟨ϵ⟩* = 0 and small variance *⟨ ϵ*^2^*⟩* = *v*_*λ*_. In a cycle with log growth factor *δ*^*d*^, the resident depletes the resource after the time *t*(*λ*_opt_, *ζ*^*d*^) of Eq. (3). Being rare, the mutant does not alter this depletion time, so a mutant cell with growth rate *λ* and lag *T* (*λ*) multiplies its biomass by 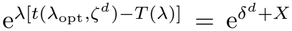, where *X* = *λ* [ *t*(*λ*_opt_, *ζ*^*d*^) *− t*(*λ, ζ*^*d*^) ] is the one-cycle advantage of the cell and *t*(*λ, ζ*^*d*^) is the time that cells with growth rate *λ* would need to complete the expansion if they were the resident. The resident multiplies its biomass by 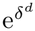, so the per-cycle change of the mutant-to-resident logabundance ratio is log *⟨*e^*X*^*⟩*, where *⟨·⟩* averages over the phenotypes of the mutant. The invasion fitness *s* is its average over cycles, and coincides with the growth rate of the mutant when rare because the log-biomass of the resident is stationary (Material and Methods). For a narrow distribution, log 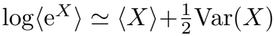, with mean and variance taken over the phenotypes of the mutant at fixed *ζ*^*d*^. Expanding 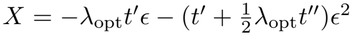 to second order in *ϵ*, where primes denote derivatives with respect to *λ* evaluated at *λ*_opt_, and using E[*t*^*′*^] = 0 gives the invasion fitness

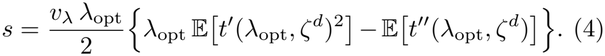

The first term is a benefit, which depends on the selection gradient *t*^*′*^ and exists only when the growth factor fluctuates. The second term is instead a cost: because *λ*_opt_ minimizes the mean completion time, cells that deviate from it in either direction are on average slower than the resident, with a penalty set by the curvature of *t*.

Since the optimal homogeneous solution minimizes the average completion time of Eq. (3), the selection gradient in cycle *d* reads

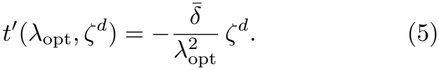

A cell with growth rate *λ*_opt_ + *ϵ* would complete the expansion earlier than the resident by about *−t*^*′*^*ϵ*, which changes its log-biomass relative to the resident by *X ≃ −λ*_opt_*t*^*′*^*ϵ*. In cycles with a large growth factor (*ζ*^*d*^ *>* 0), *t*^*′*^ *<* 0, so that faster-growing cells (*ϵ >* 0) gain biomass relative to the resident, whereas in cycles with a small growth factor (*ζ*^*d*^ *<* 0), *t*^*′*^ *>* 0, the lag weighs more and better-prepared, slower-growing cells (*ϵ <* 0) gain. This means that the sign of the selection gradient changes with the growth factor, so that different phenotypes are favored in different cycles. A heterogeneous genotype therefore always contains some cells close to the phenotype favored in the current cycle, and, because biomass grows exponentially, a cell that would complete the expansion earlier than the resident by Δ*t* gains e^*λ*Δ*t*^ *−* 1, more than the 1 *−* e^*−λ*Δ*t*^ lost by a cell that would complete it later by Δ*t*.

Explicit calculations of E[*t*^*′*2^] and E[*t*^*′′*^] (see Material and Methods) turn Eq. (4) into

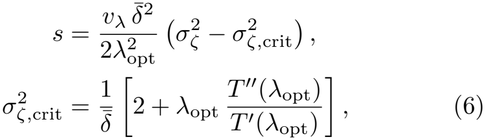

so that the invasion fitness is positive if and only if *σ*_*ζ*_ *> σ*_*ζ*,crit_. The threshold thus depends only on the mean log growth factor and on how sharply the lag increases with growth rate at the optimum.

The dependence of the lag on growth rate reflects how quickly cells rebuild the metabolic program for the incoming nutrient after a shift. Two regimes are typically observed. On the one hand, when the incoming nutrient feeds into pathways that are already active, as in shifts between glycolytic substrates (e.g., from glucose to fructose), only the entry steps have to be synthesized. Cells can devote their biosynthesis to these steps from the start, so that the missing enzymes accumulate autocatalytically, at a roughly constant rate. As a result, the lag increases only logarithmically as preparedness decreases [9]. On the other hand, shifts to gluconeogenic substrates require reversing the direction of central carbon metabolism, and the expression of gluconeogenic enzymes is itself controlled by the flux they carry [17, 25]. Poorly prepared cells therefore sustain a low flux and induce the missing enzymes slowly. This self-reinforcing bottleneck makes the lag scale inversely with preparedness, consistent with the long lags observed after shifts from glycolytic to gluconeogenic substrates [18].

These two behaviors translate into different critical thresholds. In the glycolytic case, which we use in the simulations of Figure 2, the lag is *T* = *τ*_0_ log[*f/*(1 *− f* )], with *τ*_0_ a lag time scale, and Eq. (6) takes the simple form

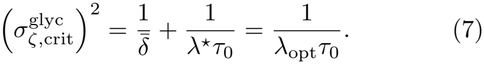

Here *λ*^*⋆*^ is the growth rate attained when the whole metabolic sector is allocated to the current nutrient. Therefore, less environmental noise is required when the mean log growth factor is large, since more of each cycle is spent growing and the optimal balance between growth and lag becomes more sensitive to the growth factor, or when lags are long compared with the growth time 1*/λ*^*⋆*^. At the same time, 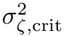 never falls below 1*/*(*λ*^*⋆*^*τ*_0_): however strong the dilution is, a minimum level of environmental noise is needed to favor heterogeneity. In the gluconeogenic case, where *T* = *τ*_0_[*f/*(1 *−f* ) *−*1], the threshold is instead

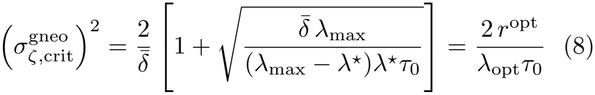

with *λ*_max_ the translation-limited maximal growth rate and *r*^opt^ = (1 *− f*_opt_)*/f*_opt_ the preparedness ratio of the resident. Now 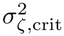 decreases as 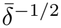 and vanishes at large mean log growth factor (see Material and Methods). When the cost of poor preparedness is this steep, even weak growth-factor fluctuations can favor phenotypic heterogeneity under strong dilution. This suggests that heterogeneity in metabolic preparedness should be selected more readily for transitions to gluconeogenic substrates, where single-cell heterogeneity has indeed been observed [25], than for transitions between glycolytic substrates.

Converted into dilution noise (see Material and Methods), the threshold *σ*_*D*,crit_ matches the onset of the favorable region in Figure 2B (dashed line); the upper edge of this region, which lies beyond the small-*σ*_*f*_ expansion, reflects the cost of broad distributions. Since the criterion depends only on *σ*_*ζ*_, supply fluctuations act like dilution fluctuations: simulations with random *c*^*d*^ confirm that adding them can push *σ*_*ζ*_ above threshold and make the heterogeneous mutant win (see Supplementary Material).

So far, we have assumed that only two prescribed strategies competed, with fixed mean allocation and different phenotypic heterogeneity. From now on, we relax such assumptions by letting both traits evolve. Specifically, at a stochastic rate, we introduce rare mutants, each changing either the mean allocation or the variability of its parent genotype, which then compete with existing genotypes under serial dilution (see Material and Methods).

We perform numerical simulations starting from different combinations of mean allocation and variability. In all evolutionary trajectories, the mean allocation first converges to *f*_opt_, irrespective of the initial variability (see Figure 3A and B). This is expected, since growth-factor fluctuations do not change the optimal homogeneous allocation. The fate of the variability, instead, depends on the dilution noise. Below threshold (see Figure 3A), the variability decreases steadily once the mean has converged, and populations become nearly homogeneous, up to a small residual variability maintained by mutation over the finite simulated time. After increasing the dilution noise above the critical threshold (see Figure 3B), trajec-tories instead settle at a finite variability, several times larger than below threshold, so that populations evolve a heterogeneous distribution of phenotypes centered on *f*_opt_. Mutants are introduced much more frequently than a sweep lasts, so that many lineages grow at the same time; the evolved variability does not depend on the supply of mutations, only the time needed to reach it does (see Supplementary Material).

**FIG 3.**
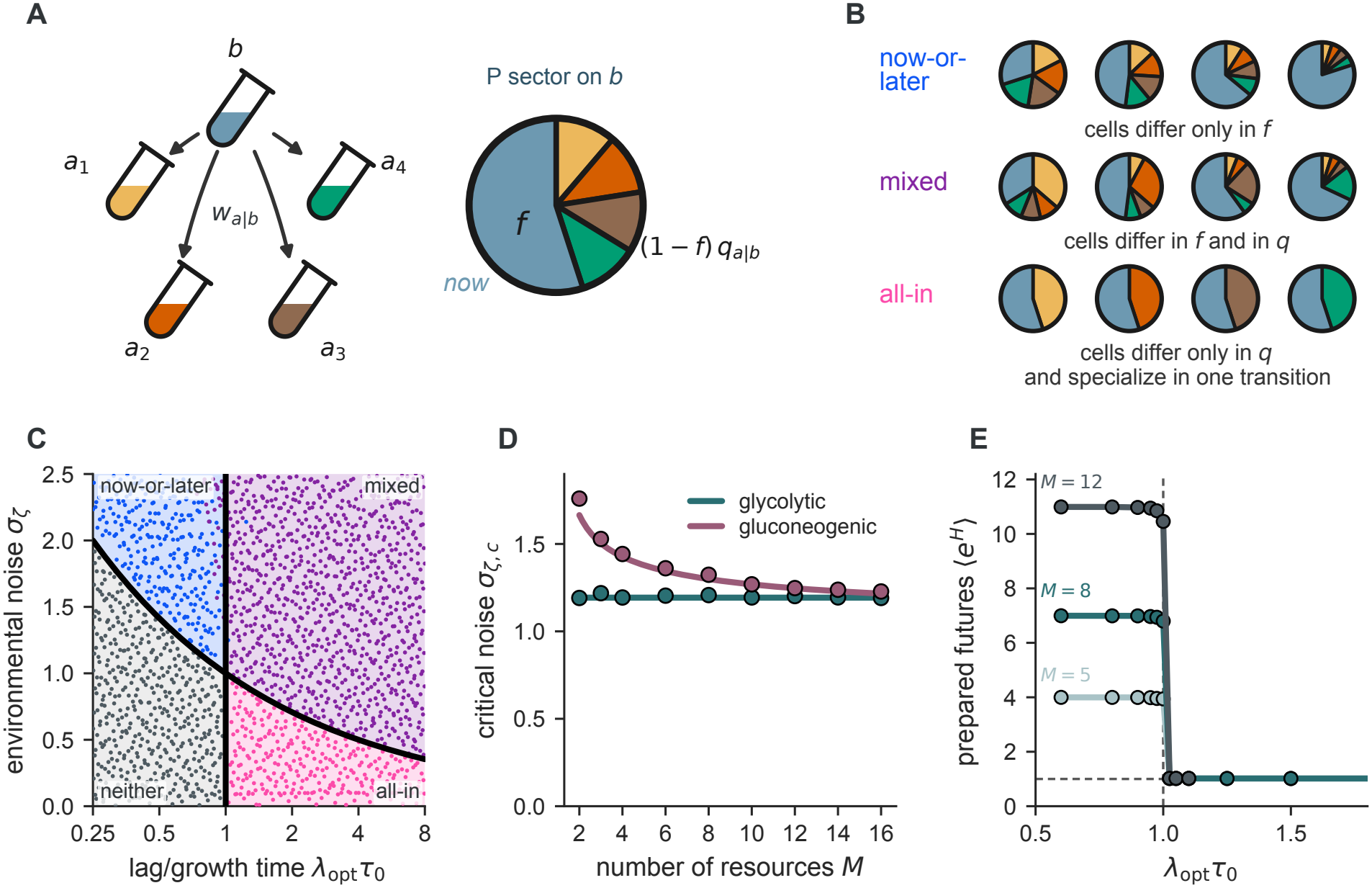
Evolution selects variability around the optimal mean allocation. (A,B) Evolutionary trajectories in the plane spanned by the biomass-weighted mean allocation 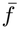 and biomass-weighted variability 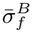. Open squares mark starting states, filled circles mark final states, and the dashed vertical line marks *f*_opt_. (C) Biomass-weighted variability 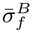 averaged over the final tail window of each run, as a function of dilution noise. Dots show independent evolutionary runs, offset horizontally for visibility, and the line shows their mean and standard error. Simulations use a constant mean log growth factor 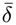; the dashed vertical line marks the corresponding local threshold converted from *σ*_*ζ*_ to *σ*_*D*_.

To summarize this behavior, we report the variability reached at the end of evolution as a function of dilution noise (see Figure 3C). Well below *σ*_*D*,crit_, the evolved variability remains at the small residual level set by mutation, independently of the noise. It starts to increase slightly before *σ*_*D*,crit_ and keeps growing with dilution noise, by more than an order of magnitude over the range explored.

Evolution thus reproduces the transition predicted by the invasion analysis and suggests cell-to-cell variability as a signature of adaptation to fluctuating environments.

### B. Many resources open a second way of hedging, which can be combined with the first

So far, a cell had to prepare for a single possible future. With *M >* 2 resources that follow one another with transition probabilities *w*_*a*|*b*_, a population experiences uncertainty not only about when the environment changes but also about which resource comes next, and this opens a second way to hedge (Figure 4A and Material and Methods). A fraction *f*_*b*_ of the metabolic sector still supports growth on the current resource *b*, while the remaining 1 *− f*_*b*_ is partitioned among the *M −* 1 alternatives,

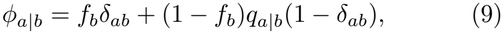

where the preparedness weight *q*_*a*|*b*_ is the share of the non-growing metabolic sector devoted to the program for *a*. For a homogeneous population and resources of equal quality, the selected weights mirror the environmental statistics, 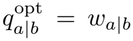 for glycolytic shifts and 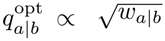 for gluconeogenic ones [9]. Both coordinates act on the lag after a shift *b → a* through a single quantity, the preparedness ratio

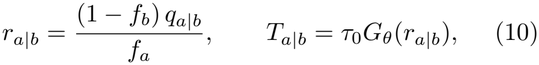

the allocation already devoted to the program for *a* relative to the one that balances growth on it, with lag set to zero for *r ≥* 1 (Material and Methods); here *G*_glyc_(*r*) = *−*log *r* for the glycolytic lag and *G*_gneo_(*r*) = 1*/r−*1 for the gluconeogenic lag, and the current allocation *f*_*a*_ also fixes the growth rate *λ*_*a*_.

**FIG 4.**
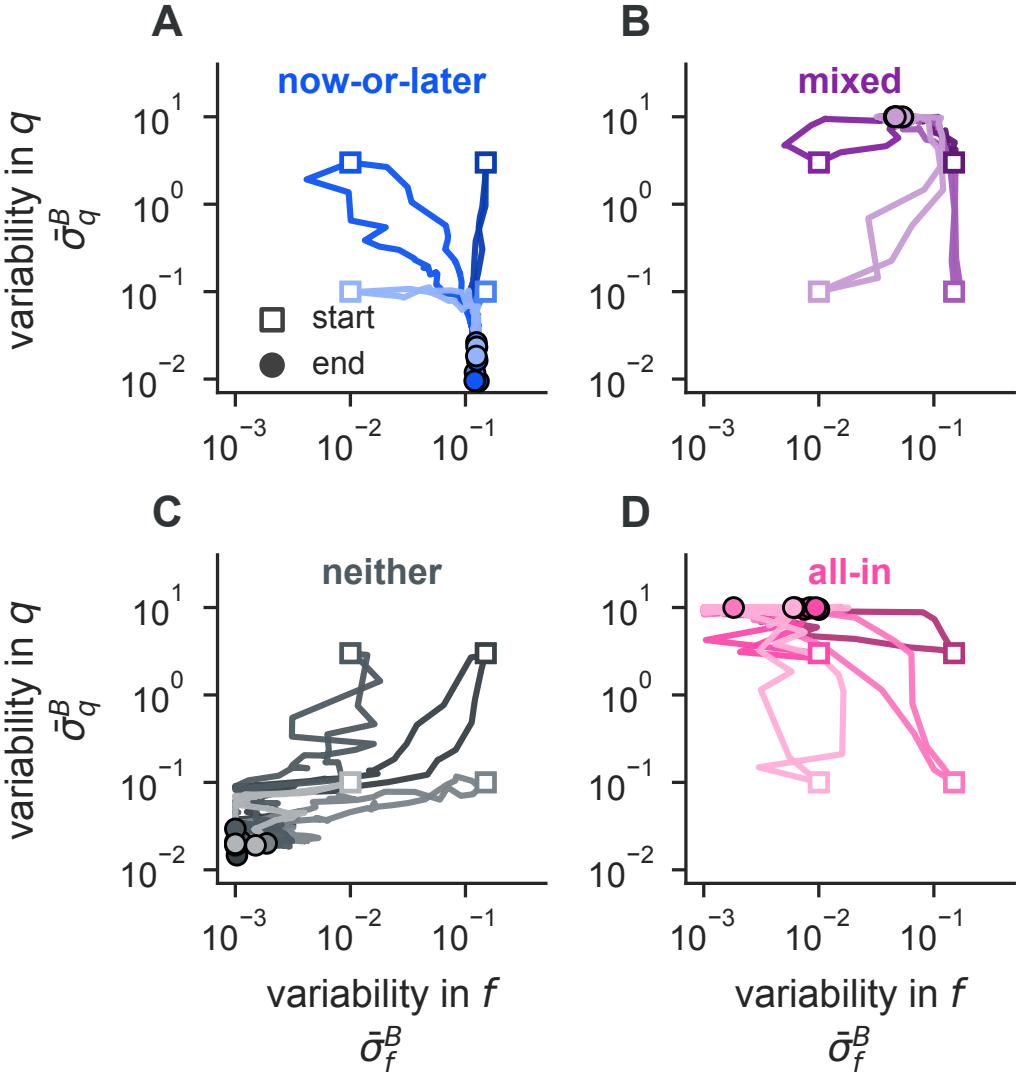
Many resources open two ways of hedging, which can be combined. (A) Resources follow one another with transition probabilities *w*_*a*|*b*_. A cell growing on *b* devotes a fraction *f* of its metabolic proteome to current growth and splits the rest among future programs with weights *q*_*a*|*b*_. (B) Alternative heterogeneous strategies. Now-or-later hedging spreads cells along *f*, each preparing evenly for all futures; all-in hedging keeps *f* fixed and spreads cells among futures, each a specialist for one of them; the mixed strategy does both. (C) Local stability of the homogeneous resident for the glycolytic lag, which describes shifts between glycolytic substrates, in the plane (*λ*_opt_*τ*_0_, *σ*_*ζ*_): homogeneous, now-or-later only, all-in only, or both are favored. Points show the winner of the competition between a homogeneous genotype and genotypes with variability in *f*, in *q* and in both. (D) Now-or-later hedging is favored for *σ*_*ζ*_ *> σ*_*ζ*,crit_. With uniform transition probabilities, *σ*_*ζ*,crit_ is independent of *M* for shifts between glycolytic substrates and decreases with *M* for shifts to gluconeogenic substrates. Lines show theory and points rare-mutant invasion estimates. (E) Effective number of prepared futures per cell, *⟨e*^*H*^ *⟩*, in evolved populations in which the mean allocation, *σ*_*f*_ and *σ*_*q*_ mutate, as a function of *λ*_opt_*τ*_0_ for the glycolytic lag at fixed *σ*_*ζ*_ *< σ*_*ζ*,crit_.

Phenotypic variability can now affect two distinct components of metabolic allocation: the fraction *f* invested in current growth and the preparedness weights *q*_*a*|*b*_ distributed among future resources. This defines three possible forms of diversification (Figure 4B). In *now-or-later* hedging, cells differ only in *f*, trading off current growth against preparedness for future conditions. In *across-future* hedging, cells share the same *f* but differ in their preparedness weights, potentially specializing in different future resources. The limiting case is *all-in* hedging, where each cell prepares for a single future resource. A *mixed* strategy combines variability in both components. Which forms of diversification are favored by selection, and whether across-future hedging evolves toward complete specialization or intermediate allocations, remain to be determined.

To see which strategy is favored, we perform an invasion analysis of a homogeneous resident with optimal allocation *f*_opt_ and weights 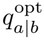, against a mutant with the same mean allocation but phenotypic variability in *f*, in *q* or in both. As for two resources, we expand the invasion fitness around the optimal homogeneous allocation, with the cells of the mutant scattered around it as *λ* = *λ*_opt_ + *ϵ* and 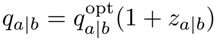 (see Material and Methods).

For the glycolytic lag and any transition matrix, aver-aging over the phenotypes and the growth factor gives the invasion fitness

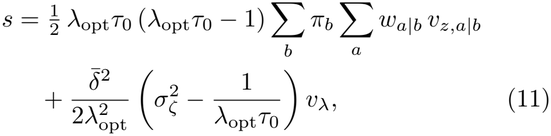

where *π*_*b*_ is the stationary distribution of the resources, 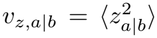 is the variance of the relative deviation of each weight, and *v*_*λ*_ = *⟨ϵ*^2^*⟩* is the variance of the growth rate, set by the variability *σ*_*f*_ of *f* . No term mixes the two strategies: the cross term is proportional to 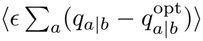 and vanishes because the weights sum to one, whatever the correlation between *ϵ* and the weights and the transition matrix. For resources of equal quality and Dirichlet distributed weights with dispersion *σ*_*q*_ around their mean, Eq. (11) simplifies to

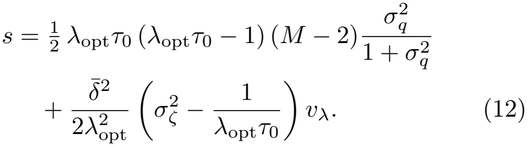

Importantly, the invasion fitness *s* is the sum of two independent terms. The first one is related to bet hedging across futures. It is present only for *M >* 2 resources, where the population can spread its preparedness weights among the alternatives. A cell whose weight on the incoming resource is *q*(1 + *z*) resumes growth with a biomass, relative to a resident cell, multiplied by the factor 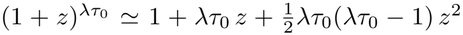. Averaged over the futures, the linear term vanishes, and the second-order term does not depend on the environment. If *λ*_opt_*τ*_0_ *>* 1 the factor is convex in the weights, so that spreading them is advantageous; at fixed mean the gain grows with *σ*_*q*_ and is largest when each cell prepares for a single future, that is, for all-in hedging. If *λ*_opt_*τ*_0_ *<* 1 the factor is concave and spreading the weights costs.

The second term is analogous to what we found for *M* = 2 resources, as it is controlled by the variability in the current al location *f* only. This term is positive only for 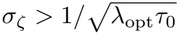, as given by Eq. (7).

Both terms are controlled by the same number, *λ*_opt_*τ*_0_, which compares the lag time to the growth time. This quantity exceeds one only when dilution is strong and the lag is longer than the growth time, since 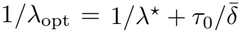.

The two conditions for the two strategies to be favored define, for the glycolytic lag, a four-region map in the plane (*λ*_opt_*τ*_0_, *σ*_*ζ*_) (see Figure 4C). When *λ*_opt_*τ*_0_ *<* 1, the first term is negative and all-in hedging is disfavored, leading to the same threshold as for two resources, 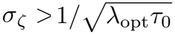, for now-or-later hedging to be favored. When *λ*_opt_*τ*_0_ *>* 1, the first term is positive and all-in hedging is favored, while both strategies are favored only if 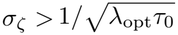. Simulations of the competition between a homogeneous genotype and genotypes with variability in *f*, in *q* or in both confirm this prediction (see points in Figure 4C).

For the gluconeogenic lag, the lag cost of a relative loss of preparedness scales as 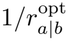 and thus depends on the future, so that the coefficients of the two terms differ among targets. For asymmetric transitions this has two consequences: the cross term between *ϵ* and *z* does not cancel after the sum over the futures, so that a covariance between the growth rate and the weights couples the two strategies, and the variance of 1*/r*^opt^ over transitions adds noise to the growth term and lowers its threshold (see Material and Methods). For symmetric transitions, however, with *w*_*a*|*b*_ = 1*/*(*M −* 1), all futures are equivalent and *r*^opt^ is the same for every transition: the covariance cancels as for the glycolytic lag, and the invasion fitness is

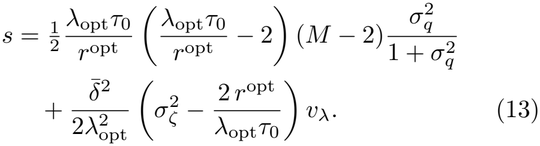

In this special case, the two strategies are again independent and the map has the same structure as for the glycolytic lag, but its thresholds depend on *r*^opt^, which in turn depends on *M* because the preparedness is divided among *M −* 1 futures; in particular the critical noise for now-or-later hedging decreases with *M* (see Figure 4D). Simulations of the invasion of rare mutants confirm this dependence. The four regions of Figure 4C also hold for the gluconeogenic lag; for asymmetric transitions the boundary of the all-in term moves to smaller values of *λ*_opt_*τ*_0_*/r*^opt^, as predicted by the local theory (see Supplementary Material).

Importantly, selection does not simply favor variability across future resources, but drives cells toward specialization. Evolutionary simulations in which the mean allocation, *σ*_*f*_, and *σ*_*q*_ mutate show that the effective number of prepared futures per cell, *⟨e*^*H*^*⟩*, where *H* is the entropy of the preparedness weights, switches from approximately *M−*1 to nearly one at *λ*_opt_*τ*_0_ = 1 (Figure 4E), rather than stabilizing at intermediate values. Thus, when diversification across futures is favored, it evolves toward an all-in strategy in which individual cells specialize in a single future resource.

To check that evolution confirms the strategy predicted in each region of Figure 4C, we let genotypes mutate in one setting per region, starting from four combinations of levels of variability (*σ*_*f*_, *σ*_*q*_) (see Figure 5). In every region, the trajectories converge to the same state, irrespective of the initial condition. In the now-or-later region *σ*_*f*_ settles well above 0, while *σ*_*q*_ decays (see Figure 5A). Where both terms are favored, variability in both *f* and *q* is selected, with *σ*_*q*_ saturating at its bound: the mixed state is reached, with a variability in *f* smaller than in the now-or-later region (see Figure 5B). Where no phenotypic bet hedging is favored, both *σ*_*f*_ and *σ*_*q*_ decay and the population becomes nearly homogeneous (see Figure 5C). Finally, in the all-in region *σ*_*q*_ grows up to a numerical bound, whereas the variability in *f* remains negligible (see Figure 5D).

**FIG 5.**
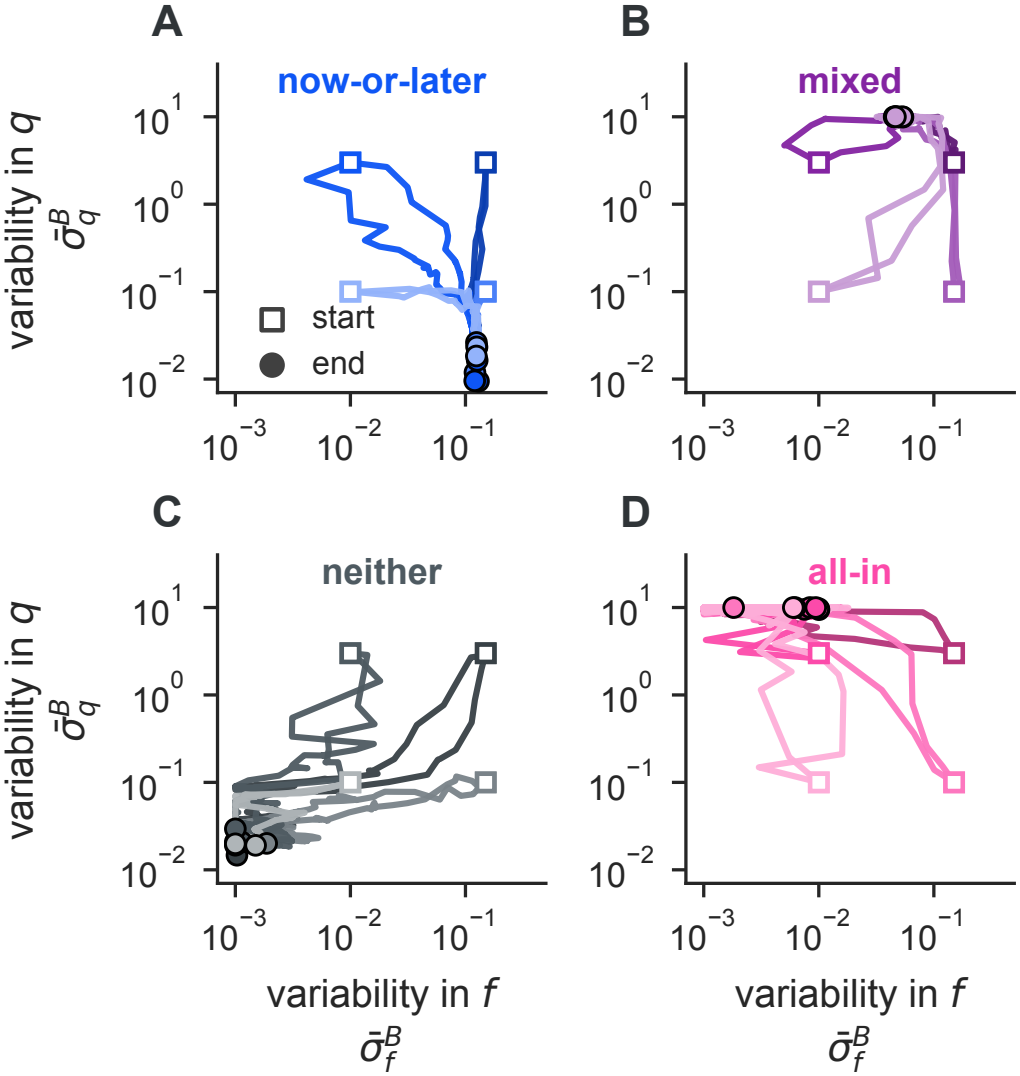
Evolution selects the strategy predicted in each region of the map. Trajectories of the biomass-weighted variability in *f* and in *q*, 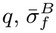 and 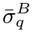, when the mean allocation, *σ*_*f*_ and *σ*_*q*_ mutate, for the glycolytic lag in one setting per region of Figure 4C, with the colors of that figure. Squares mark the four initial conditions, shown in different shades, and circles the final states, with two evolutionary runs per initial condition. Each simulation is in agreement with the predicted region.

We investigated when phenotypic heterogeneity in metabolic allocation can outperform the best homogeneous strategy in serial-dilution environments, where fluctuations in dilution or resource supply alter the growth factor of each cycle. We find that phenotypic heterogeneity is not universally advantageous, but can be selected through two distinct mechanisms. First, when fluctuations in the growth factor exceed a critical threshold, *σ*_*ζ*_ *> σ*_*ζ*,crit_, cells that strike different balances between current growth and future preparedness give an advantage to the genotype, which we call now-or-later hedging. Second, when multiple future resources are possible, having different cells prepare for different future resources can be advantageous whenever the lag is sufficiently long relative to the growth time, independently of environmental noise, which we call all-in hedging. This second route is favored whenever the lag is long compared to the growth time, *λ*_opt_*τ*_0_ *>* 1 for glycolytic lags, regardless of the noise in supply and dilution. The two conditions define four regimes, which competition and evolution simulations confirm (Figure 4C and Figure 5).

Selection acts differently on the mean and the variability of metabolic allocation. For narrow phenotypic distributions, the selected mean coincides with the optimal homogeneous allocation, which minimizes the average time required to complete a cycle and depends on the mean log growth factor, but not on its fluctuations. Environmental noise instead acts on phenotypic variability by changing the direction of selection across cycles: faster-growing cells are favored when the growth factor is high, whereas better-prepared cells gain an advantage when it is low. Because biomass grows exponentially, the gains of cells that complete the required expansion earlier can outweigh the losses of cells that complete it later by the same time. With multiple resources, the mean preparedness weights reflect the transition probabilities [9], while variability around these means hedges against uncertainty about which resource comes next. Preparation and diversification, two classic responses to environmental uncertainty [8, 10], are therefore not competing strategies, as they act on different features of the same phenotypic distribution, its mean and its variability.

The two mechanisms correspond to distinct ways of adapting to environmental uncertainty. Now-or-later hedging follows the classical evolution of phenotypic variance under fluctuating selection [27]: its benefit is proportional to the variance of the selection gradient across cycles, while its cost is determined by the curvature of fitness around the optimum. The singular variability identified in Figure 2C is the level at which the selection gradient on variability vanishes; selection converges toward it and nearby mutants cannot invade it, so it represents an evolutionarily stable level of phenotypic heterogeneity [28, 29].

All-in hedging, instead, relies on diversification among specialists. For glycolytic shifts, doubling the prepared-ness for the incoming resource shortens the lag by *τ*_0_ log 2, multiplying the relative biomass by 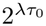. When *λτ*_0_ *>* 1, this gain is more than proportional to the investment, favoring a mixture of specialists over a generalist [4]. At large *σ*_*q*_, cells specialize in a single future resource, with the fraction of cells prepared for each resource approaching its mean preparedness weight, which for glycolytic shifts coincides with the transition probability. This resembles proportional betting in information theory and classical bet-hedging models [5, 11, 30], where populations distribute their phenotypes according to the probabilities of future environments. However, proportional betting is optimal in those models when only individuals matched to the realized environment reproduce. In our framework, even poorly prepared cells can resume growth after a finite lag, and whether proportional betting remains optimal is an open question.

Related theory has identified evolutionarily stable heterogeneity in lag when the durations of growth cycles are externally imposed [26]. Here, instead, the duration of each cycle emerges from resource depletion and population growth, with fluctuations in dilution and resource supply affecting selection through the resulting growth factor.

These predictions can be tested experimentally by vary-ing environmental noise while keeping the mean log growth factor fixed. Serial-transfer experiments with fluctuating dilution or resource supply should reveal a threshold: evolved phenotypic variability remains near its baseline below it and increases above it, whereas genotypes expressing excessively broad allocation distributions should lose in competition (Figure 2B). This adaptation may be invisible in population averages. In the two-resource model, evolution preserves the optimal homogeneous allocation as the mean allocation while increasing cell-to-cell variability (Figure 3), so populations evolved under constant and fluctuating dilution with the same mean log growth factor should differ in their allocation distributions, but not in their mean. Single-cell reporters of preparedness for alternative resources could further distinguish the two hedging strategies: now-or-later hedging predicts positive correlations in preparedness across future resources, whereas all-in hedging predicts negative correlations as cells specialize in different futures. Because lag-time heterogeneity also arises from factors unrelated to metabolic allocation [24], the relevant signature is the response of phenotypic variability to environmental noise, rather than its absolute magnitude. Lag times can evolve over the timescales of serial-transfer experiments [31], making these predictions experimentally accessible.

The critical noise threshold depends on how sharply the lag increases as metabolic preparedness decreases. For glycolytic shifts, 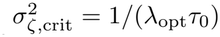, implying that, unless the lag is sufficiently long relative to the growth time, fluctuations in the growth factor must be compa-rable to its mean for now-or-later hedging to be favored. When noise arises from dilution, this requires substantial fluctuations in the logarithm of the dilution factor. For gluconeogenic shifts, where the lag increases inversely with preparedness, the threshold instead decreases with the mean log growth factor and, for symmetric resource transitions, with the number of possible future resources (Figure 4D). These shifts are also characterized by particularly long lags [18], which favor all-in hedging, and have been shown to produce bistable metabolic states [25]. Gluconeogenic shifts therefore provide a promising setting in which to experimentally investigate the evolution of heterogeneous metabolic preparedness.

Our framework relies on several simplifying assumptions that suggest directions for future work. First, we assume that allocation phenotypes are independently drawn at each cycle, neglecting phenotypic memory. In reality, metabolic states can persist across generations, and lag times depend on the history of individual lineages [32, 33]. Such memory could make selection sensitive to temporal correlations in the growth factor, as observed for stochastic phenotype switching [10, 14]. Second, we allow phenotypic variability to evolve independently of mean allocation and without an intrinsic cost, although gene expression noise is constrained by its underlying molecular mechanisms and coupled to mean expression [34, 35]. Finally, we assume that cells are transferred immediately after resource depletion, neglecting the stationary phase typical of serial-transfer experiments, which can prolong the subsequent lag [36] and introduce additional variability across cycles.

Our analytical predictions are valid for small phenotypic variability: they determine when narrow distributions can invade a homogeneous resident, but not the variability ultimately selected by evolution. The latter is constrained by the cost of broad distributions (Figure 2C), while in the many-resource simulations the degree of specialization is also limited by the maximum dispersion allowed for preparedness weights (see Supplementary Material). Consequently, the four predicted regimes accurately describe competitions at low variability, but deviations can emerge near their boundaries and at larger variability. Our analytical framework assumes that mutations are rare enough and selection is strong enough that mutation that go to fixation do so before the next one arrives (strong selection weak mutation regime [37]). Our evolutionary simulations are not restricted to that regime, as many beneficial lineages grow simultaneously and compete under clonal interference. In the Supplementary Material we show that the long-term attractor of evolutionary dynamics is independent of the population genetics regime. However, as expected [37, 38], the time to reach the evolutionary attractor (speed of adaptation) and the trajectory in phenotypic space do depend on the population genetic regime. Mapping our model with temporal and phenotypic vari-ability into know population genetic regimes, building on previous efforts [39, 40], would be necessary to describe the evolutionary trajectories of real populations.

A further open question concerns whether phenotypic diversification is favored over genetic diversity. A population can exhibit heterogeneous metabolic allocations either through different phenotypes of the same genotype or through the coexistence of distinct genotypes. In serial dilution, growth–lag trade-offs can generate frequency-dependent selection and maintain such coexistence [41, 42]. Whether environmental fluctuations favor phenotypic heterogeneity within genotypes or polymorphism among genotypes, when both strategies are available, remains to be established.

Phenotypic heterogeneity in microbes can arise as a by-product of noisy gene expression or evolve as an adaptation [43]. At the community level, conserved geneabundance distributions have been found to coexist with structured variation in gene expression in marine diatoms [44], although this pattern does not directly establish within-genotype bet hedging. Our results identify the conditions under which heterogeneity in metabolic pre-paredness becomes selectively advantageous. Fluctuations in the growth factor favor variability in how much cells invest in future resources when environmental noise exceeds a critical threshold, whereas sufficiently long lags favor having different cells prepare for different future resources. Importantly, in the two-resource model, evolution can increase phenotypic variability without changing the optimal mean allocation. Adaptation to environmental uncertainty may therefore be encoded in the distribution of phenotypes rather than in population averages.

## III. METHODS

### A. Serial-dilution ecology

We model a microbial population that repeatedly experiences growth on one limiting resource followed by dilution into fresh medium. Cycle *d* starts immediately after dilution, with genotype biomasses 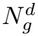. The medium contains resource *E*^*d*^ = *a* at supply 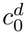. The previous resource was *E*^*d−*1^ = *b*, so the realized transition is *b→ a*. Resource identities are generated by a Markov transition matrix

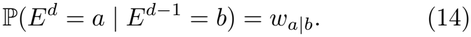

Cells grow until the supplied resource is depleted at time 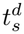, and the final biomass is divided by the dilution factor *D*^*d*^. The minimal environment used for the main theory has two resources, alternating transitions, fixed supply, and stochastic dilution. Random supply and *M >* 2 resources are introduced as extensions of the same growth– dilution map.

### B. Allocation phenotypes regulate growth and lag

The physiological state of a cell is represented by an allocation matrix ***ϕ*** over resource-specific metabolic pro-grams. The entry *ϕ*_*a*|*b*_ is the normalized fraction of the metabolic sector allocated to the program for resource *a* while the cell is prepared for resource *b*. Columns sum to one:

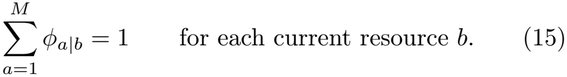

This normalized ***ϕ*** is the object used by the simulator for growth and lag. The corresponding absolute proteome mass in program *a* is *φ*_*P*_ *ϕ*_*a*|*b*_, where *φ*_*P*_ is the metabolic-sector mass fraction.

It is useful to separate the diagonal allocation from the off-diagonal preparedness pattern. For resource *b*, let *f*_*b*_ = *ϕ*_*b*|*b*_ be the fraction of the metabolic sector devoted to the current resource. The remaining fraction 1 *− f*_*b*_ is distributed among future-resource programs by normalized weights *q*_*a*|*b*_:

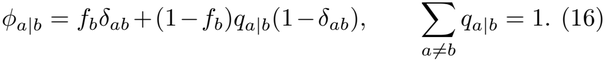

In the symmetric two-resource reduction, there is only one independent diagonal coordinate,

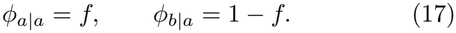

Increasing *f* improves growth on the current resource but reduces pre-existing machinery for the next resource after a shift. Thus the same coordinate controls the current-growth and future-lag sides of the trade-off.

Growth follows the standard coarse-grained proteome-allocation logic in which ribosomal and metabolic fluxes are balanced [9, 16, 45, 46]. We divide the non-housekeeping proteome into a ribosomal sector with mass fraction *φ*_*R*_ and a metabolic sector with mass fraction *φ*_*P*_ . The housekeeping sector has fixed mass fraction *φ*_*H*_ and includes the inactive ribosomes, so that *φ*_*R*_ is the active ribosomal fraction [9]. These sectors satisfy

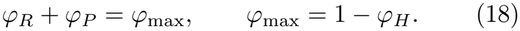

Balanced exponential growth equates the translation flux and nutrient flux:

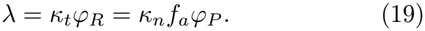

Here *κ*_*t*_ is the translation efficiency, *κ*_*n*_ is the intrinsic nutrient uptake efficiency, and only the fraction *f*_*a*_ of the metabolic sector allocated to the supplied resource contributes to uptake. For a phenotype with diagonal allocation *f*_*a*_ = *ϕ*_*a*|*a*_, the asymptotic growth rate on resource *a* is

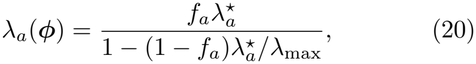

where 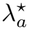 is the growth rate obtained when the full metabolic sector is allocated to resource *a*, and *λ*_max_ is the translation-limited maximum growth rate. Most numerical results use symmetric resources,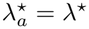.

Lag is defined as the intercept of the transient adaptation curve with the asymptotic exponential-growth trajectory. If a cell shifts from *b* to *a*, the relevant preparedness ratio is

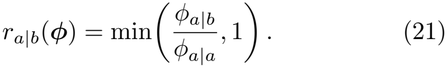

The clipping at one prevents negative lags when the preexisting allocation already exceeds the balanced-growth target. For the family of remodeling functions used here, the lag is

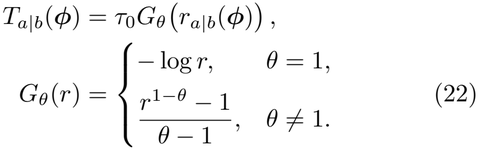

We use two members of this family throughout the paper. The logarithmic branch, appropriate for shifts between glycolytic substrates, is

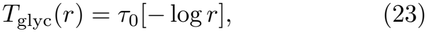

and the steeper inverse-preparedness branch, appropriate for shifts to gluconeogenic substrates, is

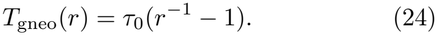

In the symmetric two-resource reduction, *r* = (1*− f* )*/f* on the positive-lag branch *f >* 1*/*2, so these become

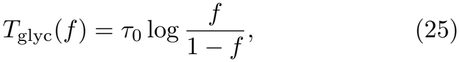

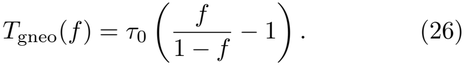

### c. Phenotypically heterogeneous genotypes

A genotype may express a single allocation phenotype or a distribution of phenotypes. We write this within-genotype phenotype density as *ρ*_*g*_(***ϕ***), normalized over the feasible allocation space. The single-phenotype case is the singular distribution *ρ*_*g*_(***ϕ***) = *δ*(***ϕ****−****ϕ***_*g*_). In the two-resource reduction, a heterogeneous genotype is summarized by a mean allocation 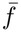 and variability *σ*_*f*_ . For *M >* 2, an additional variability can act on the off-diagonal allocation weights *q*_*a*|*b*_.

Phenotypic heterogeneity is not treated as a mixture of independently inherited lineages. At the start of a cycle, genotype biomass is distributed across the phenotypes specified by *ρ*_*g*_. Those phenotypes grow with their own growth rates and lags, but after growth and dilution their biomasses are aggregated back into the parent genotype. Selection therefore acts on total genotype biomass. This aggregation is the modeling step that distinguishes within-genotype bet hedging from coexistence among genetically distinct strategies.

For a realized transition *b → a*, define the expansion factor of phenotype ***ϕ*** over a cycle ending at time *t* as

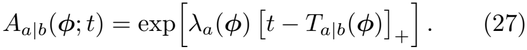

The depletion time is determined by all expressed phenotypes through

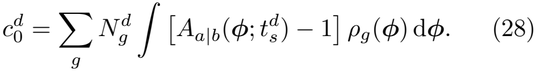

After depletion and dilution, genotype biomasses update as

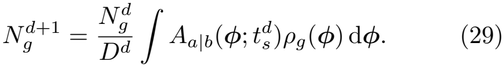

Equations (28) and (29) are the ecological core used by the simulations.

### B. Environmental noise

The environmental variable entering the local theory is the realized cycle growth factor, not the imposed physical noise by itself. For a homogeneous resident, Eq. (29) gives the logarithmic expansion the resident must achieve before dilution:

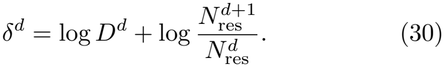

Equivalently, when the resident fully depletes the resource,

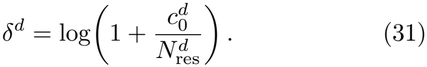

At stationarity, the long-time mean of 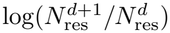 zero, so 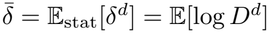. We define normalized growth-factor fluctuations by

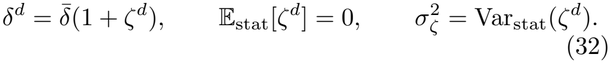

If dilution is the only fluctuating input and resident carryover is ignored, *σ*_*ζ*_ reduces to the imposed normalized dilution noise 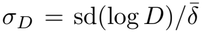. In the serial-dilution map, however, the resident biomass filters imposed dilution fluctuations, and random supply enters directly through Eq. (31). For that reason, theory is formulated in *σ*_*ζ*_. When a main-text figure uses the experimentally natural control *σ*_*D*_, the theoretical threshold is first converted through the resident biomass map.

### E. Rare invasion and local stability

Evolutionary change is modeled by rare invasion. A resident is allowed to determine the ecological depletion time, and a mutant introduced at negligible biomass invades if its invasion fitness, the long-run growth exponent in the resident environment, is positive. For resident *ρ*_res_ and mutant *ρ*_mut_, the one-cycle mutant log growth is

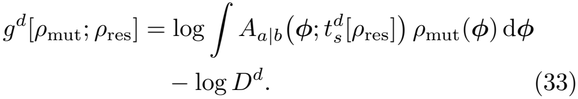

The mutant invades when

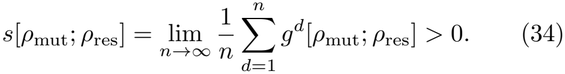

The analytical threshold is obtained for a homogeneous resident ***ϕ***_opt_ and a mutant with the same mean allocation but small phenotypic variance. For a shift *b → a* and a cycle growth factor *δ*, the completion time of a phenotype ***ϕ*** is

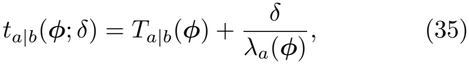

the lag paid after the shift plus the time needed to grow by *δ*. For the resident, Eqs. (29) and (30) give 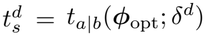 the depletion time is the completion time of the realized growth factor. As long as the phenotypes involved resume growth before depletion, so that the positive part in Eq. (27) does not bind, the mutant log growth of Eq. (33) separates into the resident growth and the one-cycle advantage *X* of the mutant phenotypes,

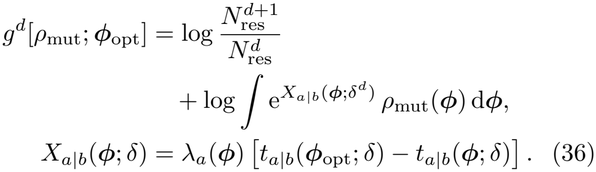

The one-cycle advantage *X* is positive when the phenotype completes the same biomass expansion sooner than the resident. The first term of *g*^*d*^ has zero mean at resident stationarity and thus does not contribute to determining the fate of the invasion. For a narrow mutant distribution the logarithm is expanded in cumulants, 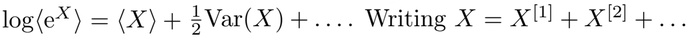 in powers of the deviation of the phenotype from the resident, with deviations of zero mean so that *⟨X*^[1]^ *⟩* = 0, the invasion fitness is where *⟨*·*⟩* averages over the phenotypes of the mutant and E over transitions and growth factors in the resident environment.

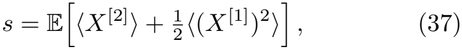

### F. Local stability with 2 resources

For two resources a phenotype is specified by *f*, and since the growth rate *λ*(*f* ) is monotone we take *λ* = *λ*_opt_ + *ϵ* as coordinate, with *⟨ϵ⟩* = 0 and 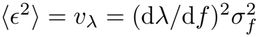, where

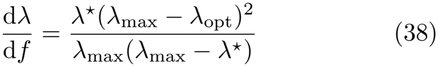

at *f*_opt_. The lag is then a function *T* (*λ*) and *t* = *T* (*λ*)+*δ/λ*. Differentiating the one-cycle advantage *X* with respect to the growth rate *λ* at the resident, with primes denoting derivatives, and using that the resident minimizes the mean completion time, E[*t*^*′*^] = 0, which holds for any variance of the growth factor because E[*ζ*^*d*^] = 0, we obtain from Eq. (37)

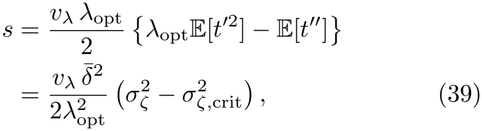

with

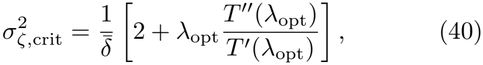

so that the homogeneous optimum is unstable to phenotypic variance if and only if *σ*_*ζ*_ *> σ*_*ζ*,crit_. In the symmetric two-resource model, explicit calculations with the lag laws of Eq. (22) lead to

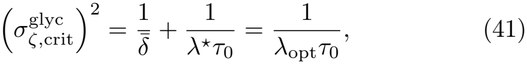

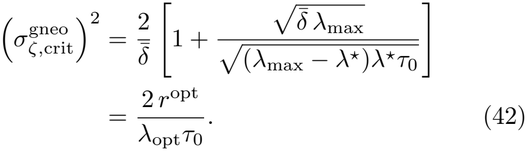

Here *r*^opt^ = (1 *− f*_opt_)*/f*_opt_ is the preparedness ratio of the resident, and the last forms follow from the condition that the resident minimizes the mean completion time (see Supplementary Material). For the glycolytic lag the threshold is thus set by the lag time in units of the growth time, *λ*_opt_*τ*_0_, and never falls below 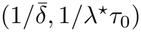.

### G. Local stability with many resources

For *M >* 2 we take resources of equal quality, pheno-types with a single diagonal allocation *f*, no self transitions, and a growth-factor distribution independent of the transition. At the resident optimum the preparedness weights are 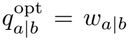 for the glycolytic lag and 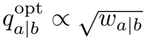 for the gluconeogenic lag [9]. A phenotype is specified by its growth rate and its weights, and for each shift *b → a* the one-cycle advantage *X*_*a*|*b*_ reads

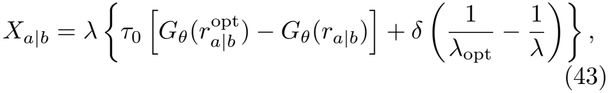

the first term being the lag saved and the second the change in the time needed to grow by *δ*. The advantage is thus a function of the growth rate *λ* and the preparedness ratio *r*, which factorizes as

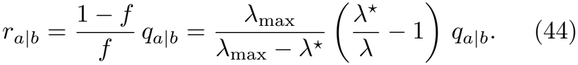

We expand the growth rate and the preparedness weights around the resident optimum as

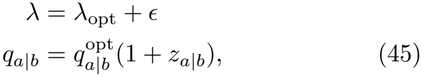

with *⟨ϵ⟩* = *⟨z⟩* = 0 and, by normalization, 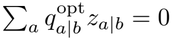.

Thus, *ϵ* sets the growth rate and the total preparedness, and *z* how the latter is divided among the futures.

Expanding the one-cycle advantage *X* to the second order gives

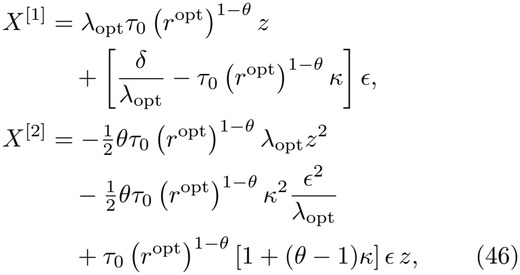

where *κ* = *λ*^*⋆*^*/*(*λ*^*⋆*^ *− λ*_opt_) = *−*d log[(1 *− f* )*/f* ]*/*d log *λ* measures how fast the preparedness budget (1 *− f* )*/f* decreases with the growth rate (see Supplementary Material), and the indices *a*|*b* of *z* and *r*^opt^ are left implicit.

#### 1. Glycolytic lag

For the glycolytic lag, *θ* = 1 and *G*_glyc_(*r*) =*−* log *r*, so that (*r*^opt^)^1*−θ*^ = 1 and Eq. (46) becomes

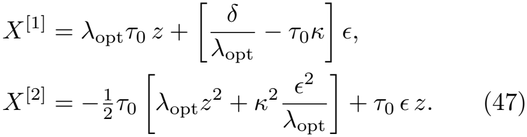

The resident minimizes the mean completion time, which for the glycolytic lag fixes 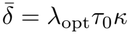 on every transition, equivalently 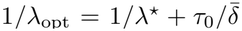. Averaging over the phenotypes and the growth factor, with 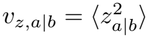 and *v*_*λ*_ = *⟨ϵ*^2^*⟩*, the two contributions to the invasion fitness are

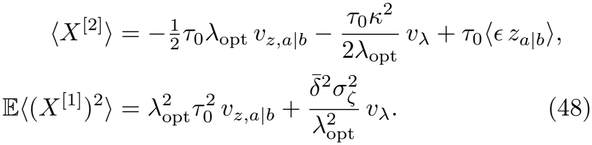

The covariance *ϵ z*_*a*|*b*_ of a single target need not vanish, but it cancels after the sum over the futures: since *q*^opt^ = *w*, the weighted deviations 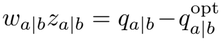 add up to zero for every phenotype, so that 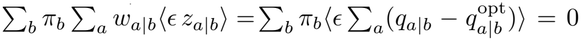, whatever the correlation between *ϵ* and the weights. Collecting the terms, and using 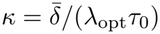,

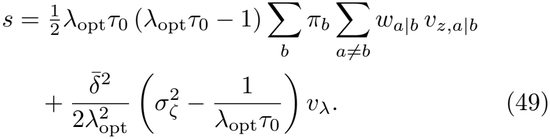

The first term is the contribution of variability in the preparedness weights, whereas the second is the contribution of variability in the growth rate, and no term mixes the two. For Dirichlet-distributed weights with concentration 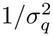, Eq. (49) simplifies to

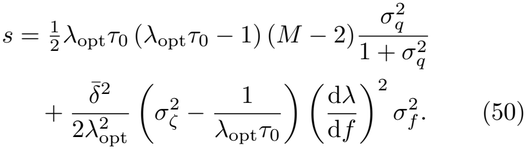

As expected, the preparedness term is absent for *M* = 2. For *M >* 2, it is positive if and only if *λ*_opt_*τ*_0_ *>* 1. On the other hand, the growth term is positive if and only if *σ*_*ζ*_ *> σ*_*ζ*,crit_ with 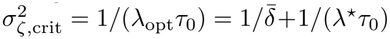. Notably, this is the same threshold of Eq. (41), which therefore does not depend on *M* : dividing preparedness among *M −* 1 alternatives adds only the constant *τ*_0_ log(*M −* 1) to the lag.

#### 2. Gluconeogenic lag

For the gluconeogenic lag, *θ* = 2 and *G*_gneo_(*r*) = 1*/r −* 1, so that 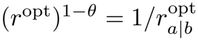. With respect to the glycolytic lag, *τ*_0_ is replaced by 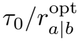 in every term of Eq. (46), so that the lag cost of a relative loss of preparedness depends on the target, the curvature terms double, and the cross term has coefficient 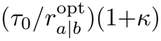. The resident minimizes the mean completion time, which fixes 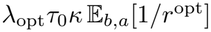, where E_*b,a*_ averages over transitions with weights *π*_*b*_*w*_*a*|*b*_. Averaging as for the glycolytic lag, for any transition matrix,

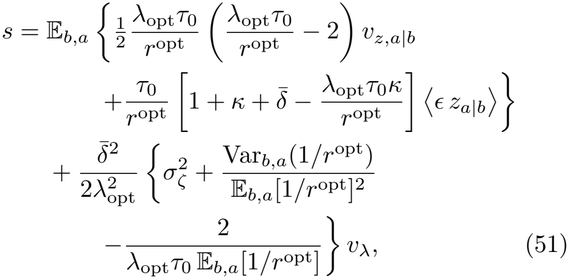

with 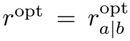 inside the averages. Compared with the glycolytic lag, the coefficient of the preparedness term depends on the target, the growth term acquires the variance of 1*/r*^opt^ over transitions, which acts as additional noise in the coefficient of *ϵ*, and the covariance term does not cancel in general, because its coefficient depends on the target also through the deviation of each transition from stationarity.

For symmetric transitions, *w*_*a*|*b*_ = 1*/*(*M* − 1) for every target, so that 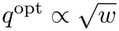 is also uniform, *q*^opt^ = 1*/*(*M* − 1), and all targets have the same *r*^opt^. The variance in Eq. (51) vanishes, stationarity holds on every transition, 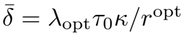, and the coefficient of the covariance is the same for every target, so that it cancels after the sum over the futures as for the glycolytic lag. For Dirichlet-distributed weights the result has the structure of Eq. (50), with *λ*_opt_*τ*_0_ replaced by *λ*_opt_*τ*_0_*/r*^opt^ and the number 1 in both brackets replaced by 2,

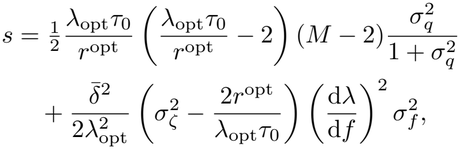

and the critical noise of the growth term, 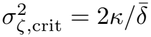, is

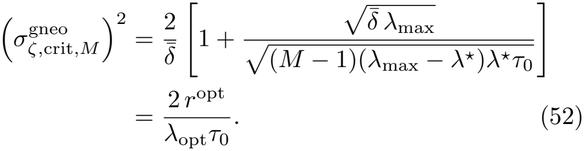

Unlike the glycolytic threshold, it decreases with *M*, because dividing preparedness among *M −*1 alternatives multiplies the allocation-dependent part of the lag by *M −* 1.

## AKNOWLEDGMENTS

We thank A. Goyal, M. Dal Bello, L. Fant, and W. Shoemaker for insightful discussions. We are particularly grateful to W. Shoemaker for the critical discussions on the population genetics aspect that are and are not included in this work. We thank Fondo Italiano per la Scienza - FIS (CUP J53C23002290001) and the Human Frontier Science Program (grant RGP004/2025) for support.

