## Supplementary Material for "Environmental Noise and Lag Select Distinct Phenotypic Bet Hedging in Microbial Metabolic Allocation"

(Dated: November 27, 2026)

### CONTENTS

I. Model Definition 2
A. Proteome sectors and allocation states 2
B. Growth–dilution dynamics for single-phenotype genotypes 2
C. Growth–dilution dynamics for phenotypically heterogeneous genotypes 4

II. Invasion Analysis 5
A. Rare mutants grow in the resident environment 5
B. Cycle growth factor in a homogeneous resident environment 6
C. Stationary environmental noise 6
D. Small phenotypic variance tests the homogeneous resident 7

III. Environmental Noise Destabilizes the Optimal Homogeneous Allocation in Two-Resource Environments 7
A. The growth-lag tradeoff determines the homogeneous optimum 8
B. Cycle-growth-factor fluctuations destabilize the homogeneous optimum 8

IV. Many Resources: Growth and Future Preparedness 11
A. The homogeneous resident defines the preparedness pattern 11
B. The local invasion criterion uses growth and completion time 12
C. The one-cycle advantage expands to second order in the growth and share deviations 12
D. The glycolytic invasion fitness is the sum of a preparedness term and a growth term 13
E. The gluconeogenic lag keeps the same two terms but its cost depends on the target 14
F. For symmetric resources the optimum and the thresholds take closed forms 16

V. Numerical Simulations 18
A. Simulation Methods 18
B. Fixed-strategy assays and evolutionary runs use the same simulator and differ only in the protocol 19
C. Environmental noise depends on dilution and supply fluctuations 20
D. Supply fluctuations enter through the cycle growth factor 22
E. The four regions of the map hold for the gluconeogenic lag and shift for non-symmetric transitions 22
F. The evolved variability does not depend on the supply of mutations 23

References 26

### I. MODEL DEFINITION

We model a microbial population that experiences repeated growth–dilution cycles. In each cycle, the medium contains a single limiting resource; cells grow until that resource is depleted, and a fraction of the final population is transferred to the next medium. The model couples two levels of description. At the physiological level, proteome allocation sets both the growth rate in the supplied resource and the lag time after a resource switch [1–3]. At the population level, a genotype may express either a single allocation phenotype or a distribution of allocation phenotypes [3, 4].

#### A. Proteome sectors and allocation states

We divide the proteome into three coarse-grained sectors: housekeeping (H), ribosomal (R), and metabolic (P). The housekeeping sector has fixed mass fraction $\varphi_H$. The ribosomal sector has mass fraction $\varphi_R$ and controls translation. The metabolic sector has mass fraction $\varphi_P$ and is responsible for mass flux from nutrients to amino acids. Here, we use $\varphi$ for proteome-sector mass fractions and $\phi$ for allocation fractions within the metabolic sector. Inactive ribosomes are counted in the housekeeping sector, as in Ref. [5], so that the active ribosomal and metabolic sectors satisfy

$$\varphi_R + \varphi_P = \varphi_{\max}, \qquad \varphi_{\max} = 1 - \varphi_H. \tag{1}$$

Under balanced exponential growth, the population growth rate $\lambda$ is set by the equality between the nutrient-to-amino-acid flux and the amino-acid-to-protein flux,

$$\lambda = \kappa_t \varphi_R = \kappa_n \varphi_P, \tag{2}$$

where $\kappa_t$ is the translation efficiency and $\kappa_n$ is the nutrient uptake efficiency. Only the metabolic allocation to the supplied resource contributes to $\kappa_n$ in that cycle. Increasing that allocation improves current growth, but leaves less proteome pre-allocated to resources that may arrive next. This is the growth–lag tradeoff controlled by the metabolic allocation strategy [2, 5, 6].

For $M$ possible resources, $\phi_{a|b}$ denotes the fraction of the metabolic sector assigned to program $a$ in cells currently adapted to resource $b$. The allocation matrix $\boldsymbol{\phi}$ has columns that sum to one. Its diagonal entry $f_b = \phi_{b|b}$ is the allocation to the current resource. The remaining fraction $1 - f_b$ is distributed across future-resource programs using normalized weights $q_{a|b}$:

$$\phi_{a|b} = f_b \delta_{ab} + (1 - f_b) q_{a|b} (1 - \delta_{ab}), \tag{3}$$

where $\delta_{ab}$ is the Kronecker delta and $\sum_{a \neq b} q_{a|b} = 1$.

Thus $\boldsymbol{\phi}$ is the key phenotypic variable: it determines both growth on the current resource and lag after resource switches. The admissible allocation space is the product of $M$ column simplices:

$$\mathcal{A}_M = \left\{ \boldsymbol{\phi} \in \mathbb{R}_+^{M \times M} : \sum_{a=1}^{M} \phi_{a|b} = 1 \text{ for all } b \right\}. \tag{4}$$

In the following, resource-specific uptake efficiencies are equal, so growth on resource $b$ is determined by the diagonal entry $f_b$ alone.

#### B. Growth–dilution dynamics for single-phenotype genotypes

The allocation space $\mathcal{A}_M$ defines the possible metabolic phenotypes available to a cell. We begin with the simplest case in which each genotype $g$ expresses a single phenotype, i.e., a single allocation matrix $\boldsymbol{\phi}_g \in \mathcal{A}_M$. All cells carrying $g$ have the same physiological response to a realized resource transition: the same growth rate on the current resource and the same lag determined by the previous resource. Equivalently, anticipating the distribution notation used below for phenotypic heterogeneity, a single-phenotype genotype is represented by the singular density

$$\rho_g(\boldsymbol{\phi}) = \delta(\boldsymbol{\phi} - \boldsymbol{\phi}_g). \tag{5}$$

We now embed this physiological description in repeated growth–dilution cycles. Each cycle $d$ begins immediately after dilution from the previous cycle, when genotype $g$ has biomass $N_g^d$. A fresh medium then supplies one limiting resource, with identity $E^d$ and amount $c_0^d$. Cells whose previous physiological state was prepared for the resource in the preceding cycle may first experience a lag; after that lag they grow exponentially with the growth rate set by their allocation to the supplied resource. Growth stops when the limiting resource is depleted at time $t_s^d$. The resulting biomass is then diluted by a factor $D^d$, giving the initial biomass for the next cycle. Thus selection acts on the whole cycle, through the combination of lag, exponential growth, resource depletion, and dilution.

For a realized cycle, growth depends on the supplied resource, whereas lag depends on the resource for which cells were prepared in the previous cycle. The resource identities $E^d \in \{1, \ldots, M\}$ form a Markov chain, with transition probabilities

$$\mathbb{P}(E^d = a \mid E^{d-1} = b) = w_{a|b}, \qquad \sum_{a=1}^{M} w_{a|b} = 1, \tag{6}$$

where we use the shorthand $a = E^d$ and $b = E^{d-1}$ for the current and previous resource identities in cycle $d$, respectively. The general Markov chain may include self transitions. The normalized preparedness coordinates $q_{a|b}$ in Eq. (3), however, describe only off-diagonal preparation for resource switches. When a self transition occurs, no remodeling lag is incurred in the $q$ block.

Within a cycle, let $n_g(t)$ denote the biomass of genotype $g$ at time $t$ after transfer to the fresh medium. At the continuous physiological level,

$$\frac{1}{n_g(t)} \frac{\mathrm{d}n_g(t)}{\mathrm{d}t} = \lambda_{a|b}(t; \boldsymbol{\phi}_g), \tag{7}$$

where $\lambda_{a|b}(t; \boldsymbol{\phi}_g)$ is the instantaneous growth rate during adaptation to resource $a$ after previously being in resource $b$. This time dependence comes from proteome remodeling. Let $\tilde{\phi}_{a|b}(t)$ be the normalized allocation at time $t$ to the program for the new resource $a$ during this remodeling process. Its initial value is the pre-existing allocation inherited from the previous physiological state, $\tilde{\phi}_{a|b}(0) = \phi_{a|b}$, and its asymptotic value is the balanced-growth allocation on the new resource, $\tilde{\phi}_{a|b}(\infty) = \phi_{a|a}$. At each time, the instantaneous growth rate is the flux-balance growth rate associated with the current value of this allocation,

$$\lambda_{a|b}(t; \boldsymbol{\phi}_g) = \Lambda_a\left(\tilde{\phi}_{a|b}(t)\right), \tag{8}$$

where the explicit form of $\Lambda_a$ is derived below. Thus, the biomass equation is coupled to the allocation-remodeling dynamics. We model remodeling as a saturating relaxation toward the balanced-growth allocation on the new resource. The relaxation rate is proportional to the target allocation $\phi_{a|a}$, vanishes when the target is reached, and may depend on the amount of new-resource machinery already present through a monotone factor $h_\theta$ [5]. This gives the remodeling equation

$$\frac{\mathrm{d}\tilde{\phi}_{a|b}}{\mathrm{d}t} = \frac{\phi_{a|a}}{\tau_0} \left(1 - \frac{\tilde{\phi}_{a|b}}{\phi_{a|a}}\right) h_\theta\left(\frac{\tilde{\phi}_{a|b}}{\phi_{a|a}}\right), \tag{9}$$

Here, $\tau_0$ is the characteristic time scale for the remodeling process, and $h_\theta(z) = z^\theta$ is a power-law function that captures the dependence of remodeling on the preparedness ratio $z = \tilde{\phi}_{a|b}/\phi_{a|a}$. The exponent $\theta$ controls how strongly remodeling is limited by the new-resource machinery that is already present at the moment of the switch. For $\theta = 1$, enzyme production is autocatalytic in the simplest sense: when the new program is rare, its increase is proportional to its current abundance. Metabolically, this corresponds to a switch in which the nutrient flux generated by the new enzymes limits the production of those same enzymes, while nutrient and ribosomal fluxes remain matched. For $\theta = 2$, this dependence is sharper. When the new program is rare, remodeling is quadratically suppressed, as expected for switches in which activating the new pathway requires a stronger pre-existing flux through that pathway. A relevant example is a switch from glycolytic to gluconeogenic carbon sources, where the machinery needed to produce biomass precursors through gluconeogenesis is initially scarce [2, 3].

Rather than carrying the full transient function $\lambda_{a|b}(t; \boldsymbol{\phi}_g)$ in the ecological dynamics, we use its long-time exponential approximation. The asymptotic growth rate on resource $a$ is denoted by $\lambda_a(\boldsymbol{\phi}_g)$, and the lag time is defined as the intercept of the asymptotic exponential trajectory:

$$\int_0^t \lambda_{a|b}(s; \boldsymbol{\phi}_g)\, \mathrm{d}s = \lambda_a(\boldsymbol{\phi}_g) \left[t - T_{a|b}(\boldsymbol{\phi}_g)\right] + o(1) \qquad (t \to \infty). \tag{10}$$

In this way, the full transient growth curve is replaced by no effective growth for a time $T_{a|b}$, followed by exponential growth at rate $\lambda_a$.

For growth on resource $a$, only the metabolic program for that same resource contributes to nutrient uptake. It is therefore useful to denote the relevant diagonal entry of genotype $g$ by $f_{a,g} \equiv (\boldsymbol{\phi}_g)_{a|a}$. Eliminating the ribosomal and metabolic sector sizes from Eqs. (1) and (2) gives the scalar tradeoff $\lambda_a(\boldsymbol{\phi}_g) = \Lambda_a(f_{a,g})$ with

$$\Lambda_a(f) = \frac{f\lambda_a^\star}{1-(1-f)\lambda_a^\star/\lambda_{\max}}. \tag{11}$$

Here, $\lambda_a^\star$ denotes the growth rate achieved at $f = 1$, when the entire metabolic sector is allocated to the supplied resource $a$. In the symmetric-resource regime considered here, this value is the same for all resources. The parameter $\lambda_{\max}$ denotes the translation-limited maximum growth rate.

The lag depends on how much of the new-resource program was already present before the switch. For a switch from $b$ to $a$, we assume $0 < \phi_{a|b} < \phi_{a|a}$: cells prepared for resource $b$ carry some machinery for resource $a$, but less than the balanced-growth amount expressed when $a$ is the supplied resource. If the pre-existing allocation equals or exceeds that target, the effective lag is zero rather than negative. Solving Eq. (9) and applying the intercept definition in Eq. (10) gives, for this positive-lag branch,

$$T_{a|b}(\boldsymbol{\phi}) = \tau_0 \frac{\left(\frac{\phi_{a|b}}{\phi_{a|a}}\right)^{1-\theta} - 1}{\theta - 1}, \qquad \theta \neq 1. \tag{12}$$

The logarithmic case $\theta = 1$ is obtained by taking the continuous limit $\theta \to 1$, which gives $-\tau_0 \log(\phi_{a|b}/\phi_{a|a})$. For $\theta = 2$, the lag becomes the stronger inverse-preparedness form $\tau_0(\phi_{a|a}/\phi_{a|b} - 1)$.

Once $\lambda_a$ and $T_{a|b}$ are fixed by the allocation matrix, the duration of a growth cycle is determined by resource depletion. For single-phenotype genotypes, the depletion time solves

$$c_0^d = \sum_g N_g^d \left\{ \exp\Big[\lambda_a(\boldsymbol{\phi}_g)\left[t_s^d - T_{a|b}(\boldsymbol{\phi}_g)\right]_+\Big] - 1 \right\}, \tag{13}$$

where we choose biomass units in which the yield from resource to biomass is one and use $[x]_+ = \max(x, 0)$, so that lag delays growth without causing biomass decrease. In predictable environments, this depletion time is the cycle-level object selected by the environment: the evolutionarily stable homogeneous strategy is obtained by minimizing the depletion time averaged over the environmental process, as in the monomorphic allocation theory of Ref. [5]. After depletion and dilution, genotype biomasses update deterministically as

$$N_g^{d+1} = \frac{N_g^d}{D^d} \exp\Big[\lambda_a(\boldsymbol{\phi}_g)\left[t_s^d - T_{a|b}(\boldsymbol{\phi}_g)\right]_+\Big]. \tag{14}$$

This cycle map applies to any realized environmental sequence: resource identities are generated by Eq. (6), while $c_0^d$ and $D^d$ specify the resource supply and dilution factor in cycle $d$.

#### C. Growth–dilution dynamics for phenotypically heterogeneous genotypes

So far, we have described the model in the simplest case in which each genotype expresses a single allocation phenotype. More generally, a genotype may distribute its cells across a range of allocation phenotypes. We consider this phenotypic variability as within-genotype variability: cells share the same heritable parameters, but need not express the same allocation matrix in a given cycle. We will show that such variability can be favored by selection when environmental fluctuations are sufficiently strong, because it allows a genotype to spread its biomass across phenotypes that differ in their growth–lag tradeoff and are favored in different environmental realizations.

Let $\rho_g(\boldsymbol{\phi})$ be the phenotype density produced by genotype $g$ on $\mathcal{A}_M$:

$$\rho_g(\boldsymbol{\phi}) \geq 0, \qquad \int_{\mathcal{A}_M} \rho_g(\boldsymbol{\phi})\, \mathrm{d}\boldsymbol{\phi} = 1. \tag{15}$$

Here $\mathrm{d}\boldsymbol{\phi}$ denotes integration over the independent coordinates of the product of column simplices in Eq. (4). The single-phenotype model above is recovered by the singular density in Eq. (5), with the delta understood on $\mathcal{A}_M$. At the beginning of a cycle, the genotype biomass is distributed over phenotypes according to $\rho_g$.

Equations (13) and (14) were written for a genotype represented by a single allocation matrix $\boldsymbol{\phi}_g$. For a heterogeneous genotype, each phenotype $\boldsymbol{\phi}$ in the support of $\rho_g$ follows the same physiological rules: its growth rate is $\lambda_a(\boldsymbol{\phi})$, as defined by Eq. (11), and its lag after the realized transition $b \to a$ is $T_{a|b}(\boldsymbol{\phi})$, as defined by Eq. (12). Thus the single-phenotype factors in Eqs. (13) and (14) are replaced by their averages over the phenotype density. The depletion condition becomes

$$c_0^d = \sum_g N_g^d \int_{\mathcal{A}_M} \left\{ \exp\Big[ \lambda_a(\boldsymbol{\phi}) \left[ t_s^d - T_{a|b}(\boldsymbol{\phi}) \right]_+ \Big] - 1 \right\} \rho_g(\boldsymbol{\phi}) \, \mathrm{d}\boldsymbol{\phi}. \tag{16}$$

After depletion and dilution, genotype biomasses update as

$$N_g^{d+1} = \frac{N_g^d}{D^d} \int_{\mathcal{A}_M} \exp\Big[ \lambda_a(\boldsymbol{\phi}) \left[ t_s^d - T_{a|b}(\boldsymbol{\phi}) \right]_+ \Big] \rho_g(\boldsymbol{\phi}) \, \mathrm{d}\boldsymbol{\phi}. \tag{17}$$

Equations (16) and (17) are the ecological core of the heterogeneous model. For the singular density in Eq. (5), they reduce to the single-phenotype dynamics in Eqs. (13) and (14). All evolutionary calculations below use the general equations, with different choices of phenotype density $\rho_g$, environmental process, and mutation kernel.

### II. INVASION ANALYSIS

We first formulate evolution in the rare-mutant limit. A resident genotype is first allowed to establish its ecological dynamics. A mutant genotype is then introduced at negligible biomass. The mutant experiences the environmental process generated by the resident, but does not affect the resident depletion time and biomass. The mutant invades if its invasion fitness, the long-run growth exponent in the resident environment, is positive. This separation between ecological equilibration and mutant growth gives a local stability criterion for the resident phenotype. In the main text, the same criterion is compared with fixed-strategy competitions and with explicit evolutionary simulations in which mutants are generated sequentially over many serial-dilution cycles; the simulator is described in Sec. V. The criterion describes the successional regime, in which each mutant is resolved before the next one arrives; the evolutionary simulations introduce mutants much more frequently than the duration of a sweep, and Sec. V F shows that this does not change the evolved state. Here we derive the invasion criterion for a general resident and mutant genotype, and then specialize to the case of a homogeneous resident and a mutant with small phenotypic variance.

#### A. Rare mutants grow in the resident environment

Let the resident phenotype density be $\rho_{\mathrm{res}}$ and the mutant phenotype density be $\rho_{\mathrm{mut}}$. In cycle $d$, the resident determines $t_s^d[\rho_{\mathrm{res}}]$ through Eq. (16). The mutant biomass obeys

$$N_{\mathrm{mut}}^{d+1} = \frac{N_{\mathrm{mut}}^d}{D^d} \int_{\mathcal{A}_M} \exp\Big[ \lambda_a(\boldsymbol{\phi}) \left[ t_s^d[\rho_{\mathrm{res}}] - T_{a|b}(\boldsymbol{\phi}) \right]_+ \Big] \rho_{\mathrm{mut}}(\boldsymbol{\phi}) \, \mathrm{d}\boldsymbol{\phi}, \tag{18}$$

where $a = E^d$ and $b = E^{d-1}$. Importantly, the depletion time $t_s^d$ is determined by the resident and is unaffected by the mutant, which is assumed to be at negligible biomass. The corresponding logarithmic growth increment is defined as

$$g^d[\rho_{\mathrm{mut}}; \rho_{\mathrm{res}}] = \log \int_{\mathcal{A}_M} \exp\Big[ \lambda_a(\boldsymbol{\phi}) \left[ t_s^d[\rho_{\mathrm{res}}] - T_{a|b}(\boldsymbol{\phi}) \right]_+ \Big] \rho_{\mathrm{mut}}(\boldsymbol{\phi}) \, \mathrm{d}\boldsymbol{\phi} - \log D^d. \tag{19}$$

The mutant invades if its invasion fitness, the long-run mean of the logarithmic growth increment, is positive, namely if

$$s[\rho_{\mathrm{mut}}; \rho_{\mathrm{res}}] = \lim_{n\to\infty} \frac{1}{n} \sum_{d=1}^{n} g^d[\rho_{\mathrm{mut}}; \rho_{\mathrm{res}}] > 0. \tag{20}$$

This invasion criterion is exact for the deterministic ecological dynamics in Eqs. (16) and (17).

#### B. Cycle growth factor in a homogeneous resident environment

The invasion criterion becomes explicit when the resident is homogeneous, $\rho_{\rm res}(\boldsymbol{\phi}) = \delta(\boldsymbol{\phi} - \boldsymbol{\phi}_r)$. We work in the local regime in which the relevant resident and mutant phenotypes begin growing before depletion, so the positive part does not bind. For the homogeneous resident, Eq. (17) gives

$$\log \frac{N_{\rm res}^{d+1}}{N_{\rm res}^{d}} = \lambda_a(\boldsymbol{\phi}_r)\left[t_s^d - T_{a|b}(\boldsymbol{\phi}_r)\right] - \log D^d. \tag{21}$$

It is useful to rewrite this equation in terms of the logarithmic expansion that the resident must produce during cycle $d$:

$$\delta^d = \log D^d + \log \frac{N_{\rm res}^{d+1}}{N_{\rm res}^{d}}. \tag{22}$$

For any phenotype $\boldsymbol{\phi}$ and any prescribed value of the growth factor, define the completion time

$$t_{a|b}(\boldsymbol{\phi};\delta) = T_{a|b}(\boldsymbol{\phi}) + \frac{\delta}{\lambda_a(\boldsymbol{\phi})}. \tag{23}$$

This is the lag paid after the transition plus the growth time needed to produce the logarithmic expansion $\delta$. For the homogeneous resident, Eq. (21) is equivalently

$$t_s^d = t_{a|b}(\boldsymbol{\phi}_r;\delta^d) = T_{a|b}(\boldsymbol{\phi}_r) + \frac{\delta^d}{\lambda_a(\boldsymbol{\phi}_r)}. \tag{24}$$

Thus the resident depletion time is the completion time of the resident phenotype for the realized growth factor. Mutant phenotypes are compared in the same resident environment by holding fixed the transition $b \to a$ and the growth factor $\delta^d$, while changing the allocation-dependent lag and growth.

Substituting Eq. (24) into Eq. (19) gives

$$g^d[\rho_{\rm mut};\boldsymbol{\phi}_r] = \log \frac{N_{\rm res}^{d+1}}{N_{\rm res}^{d}} + \log \int_{\mathcal{A}_M} \exp\left[X_{a|b}(\boldsymbol{\phi};\boldsymbol{\phi}_r,\delta^d)\right] \rho_{\rm mut}(\boldsymbol{\phi})\, \mathrm{d}\boldsymbol{\phi}, \tag{25}$$

where

$$X_{a|b}(\boldsymbol{\phi};\boldsymbol{\phi}_r,\delta) = \lambda_a(\boldsymbol{\phi})\left[t_{a|b}(\boldsymbol{\phi}_r;\delta) - t_{a|b}(\boldsymbol{\phi};\delta)\right]. \tag{26}$$

This is the one-cycle log-growth advantage of phenotype $\boldsymbol{\phi}$ relative to the resident environment. If $\boldsymbol{\phi}$ can satisfy the same expansion faster than the resident, then the bracket is positive and the phenotype gains biomass during that cycle.

The first term in Eq. (25) has zero long-run mean at resident stationarity. The invasion fitness of a heterogeneous mutant invading a homogeneous resident is therefore

$$s_{\boldsymbol{\phi}_r}[\rho_{\rm mut}] = \sum_b \pi_b \sum_a w_{a|b} \mathbb{E}_{\delta|a,b}\left[\log \int_{\mathcal{A}_M} \exp\left[X_{a|b}(\boldsymbol{\phi};\boldsymbol{\phi}_r,\delta)\right] \rho_{\rm mut}(\boldsymbol{\phi})\, \mathrm{d}\boldsymbol{\phi}\right], \tag{27}$$

where $\pi_b$ is the stationary probability of resource $b$. The conditional expectation is over the stationary realized growth factor generated by the resident process after the transition $b \to a$. If the mutant is also homogeneous, $\rho_{\rm mut}$ is a delta distribution and the logarithm disappears. Self transitions, when present, are included in Eq. (27) through the same Markov sum. They carry their own one-cycle growth term but do not introduce an off-diagonal preparedness coordinate.

#### C. Stationary environmental noise

The exact invasion criterion above is written in terms of the realized growth factor $\delta^d$. For local stability calculations, the useful statistical coordinate is the stationary fluctuation of the growth factor around its mean. At resident stationarity,

$$\mathbb{E}_{\rm stat}\left[\log \frac{N_{\rm res}^{d+1}}{N_{\rm res}^{d}}\right] = 0, \tag{28}$$

and Eq. (22) gives

$$\bar{\delta} = \mathbb{E}_{\text{stat}}[\delta^d] = \mathbb{E}[\log D^d]. \tag{29}$$

We then define the normalized growth-factor fluctuation by

$$\zeta^d = \frac{\delta^d - \bar{\delta}}{\bar{\delta}}, \qquad \delta^d = \bar{\delta}(1+\zeta^d), \qquad \mathbb{E}_{\text{stat}}[\zeta^d] = 0. \tag{30}$$

The environmental-noise parameter entering the local theory is the variance of this dimensionless quantity,

$$\sigma_\zeta^2 = \text{Var}_{\text{stat}}(\zeta^d) = \frac{\text{Var}_{\text{stat}}(\delta^d)}{\bar{\delta}^2}. \tag{31}$$

Thus $\sigma_\zeta$ is a property of the stationary resident environment, not of the imposed physical noise amplitudes alone. In the simplest formulation with an imposed growth factor this coordinate can be set directly; in the full biomass map, dilution and supply are filtered through resident carry-over. Its measurement and its conversion from the physical noise parameters used in simulations are described in Sec. V C.

#### D. Small phenotypic variance tests the homogeneous resident

Equation (27) is the exact invasion criterion for any heterogeneous mutant, but it does not yet say when heterogeneity itself is favored. The local evolutionary question is sharper: starting from a homogeneous resident, does a mutant genotype gain by spreading its cells over nearby allocation phenotypes while keeping approximately the same mean allocation? This is the infinitesimal version of evolutionary branching in phenotype space.

To isolate this effect, consider a mutant distribution concentrated near the resident phenotype. The logarithm in Eq. (27) can then be expanded in cumulants of $X_{a|b}$ under $\rho_{\text{mut}}$:

$$\log \int_{\mathcal{A}_M} e^{X_{a|b}(\boldsymbol{\phi})} \rho_{\text{mut}}(\boldsymbol{\phi})\, \mathrm{d}\boldsymbol{\phi} = \left\langle X_{a|b} \right\rangle_{\rho_{\text{mut}}} + \frac{1}{2}\text{Var}_{\rho_{\text{mut}}}(X_{a|b}) + O(\kappa_3). \tag{32}$$

The first cumulant is the effect of the mutant mean phenotype. The second cumulant is the leading effect of within-genotype phenotypic variance. At a monomorphic evolutionary optimum, the first-order mean shift vanishes after averaging over the resident environment. The remaining terms ask whether phenotypic variance increases or decreases the invasion fitness.

The result has a direct interpretation. Variance is costly because phenotypes away from the resident optimum experience the stabilizing curvature of the mean depletion time. At the same time, variance can also be beneficial because growth-factor fluctuations make the instantaneous selection gradient fluctuate from one cycle to the next. If this fluctuation term is larger than the curvature cost, a narrow heterogeneous mutant invades the homogeneous resident.

The next section evaluates this local condition explicitly in the symmetric two-resource reduction and shows that it can be satisfied in a regime of strong growth-factor fluctuations.

### III. ENVIRONMENTAL NOISE DESTABILIZES THE OPTIMAL HOMOGENEOUS ALLOCATION IN TWO-RESOURCE ENVIRONMENTS

The previous section reduced rare invasion to allocation-dependent growth and lag evaluated at a prescribed resident-generated cycle growth factor. We now make the local stability criterion explicit in the simplest nontrivial setting: two symmetric resource states. The environment alternates between the two resources, so each cycle begins after a switch from the other resource. We assume that the stationary distribution of the normalized fluctuation $\zeta^d$ of the growth factor is the same for the two alternating transitions.

With two symmetric resources, a homogeneous phenotype is described by one number, the allocation $f$ to the current resource:

$$\boldsymbol{\phi}(f) = \begin{pmatrix} f & 1-f \\ 1-f & f \end{pmatrix}. \tag{33}$$

The diagonal entries are the current-resource allocation, whilst the off-diagonal entries are the pre-allocation to the only possible future resource. The feasible symmetric domain is $1/2 \leq f \leq 1$: at $f = 1/2$ cells allocate equally to the two resources, while at $f = 1$ cells maximize current growth but carry no preparation for the other resource.

A heterogeneous genotype is a phenotype density $\rho(f)$ on the feasible interval. We denote its mean by $\bar{f}$ and its phenotypic standard deviation by $\sigma_f$. The homogeneous case is $\rho(f) = \delta(f - \bar{f})$, equivalently $\sigma_f = 0$.

#### A. The growth-lag tradeoff determines the homogeneous optimum

In the two-resource reduction, the same scalar $f$ controls both parts of the growth–lag tradeoff. During growth on the current resource, the diagonal allocation is $\phi_{a|a} = f$ and the remaining metabolic allocation $1 - f$ is invested in the other resource program. Equation (11) therefore becomes

$$\lambda(f) = \frac{f\lambda^\star}{1 - (1 - f)\lambda^\star/\lambda_{\max}}. \tag{34}$$

This is an increasing function of $f$: allocating more of the metabolic sector to the present resource increases the instantaneous growth rate. The same choice reduces preparation for the next resource, because in the alternating two-resource environment the incoming resource is precisely the off-diagonal program. At the switch, the amount already allocated to the incoming resource is $1 - f$, while the post-adaptation target is $f$. The preparedness ratio is therefore $(1 - f)/f$, and Eq. (12) reduces to

$$T(f) = \tau_0 G_\theta\left(\frac{1 - f}{f}\right). \tag{35}$$

Here $G_\theta(x) = (x^{1-\theta} - 1)/(\theta - 1)$ for $\theta \neq 1$, with the continuous limit $G_1(x) = -\log x$ [5]. Thus increasing $f$ creates the already mentioned physiological tension: it improves growth during the present cycle, but increases the lag paid after the next environmental switch. Selection acts on the whole depletion cycle rather than on either component separately.

For a homogeneous phenotype with allocation $f$, the completion time of Eq. (23) becomes

$$t(f, \zeta) = T(f) + \frac{\bar{\delta}(1 + \zeta)}{\lambda(f)}. \tag{36}$$

This expression separates the two contributions to cycle duration. The first term is the physiological delay after the switch, while the second one is the growth time required to produce the realized logarithmic growth factor $\bar{\delta}(1 + \zeta)$.

The optimal homogeneous allocation minimizes the environmental average of this cycle time:

$$f_{\text{opt}} = \underset{1/2 \leq f \leq 1}{\arg\min}\, \mathbb{E}_\zeta \left[t(f, \zeta)\right]. \tag{37}$$

Because $\mathbb{E}[\zeta] = 0$, the homogeneous optimum depends on the mean log growth factor $\bar{\delta}$ but not on the variance of $\zeta$. Thus the optimal homogeneous allocation is the same as in the deterministic case $\zeta = 0$: environmental fluctuations do not shift the best mean allocation of a single-phenotype genotype, as in the baseline pre-allocation problem [5], but they can change the stability of that optimum to within-genotype phenotypic variability. We therefore use $f_{\text{opt}}$ as the reference mean around which to test whether a narrow distribution of phenotypes can invade.

It is useful to write the first-order condition in terms of the growth rate $\lambda$. Let $f(\lambda)$ be the inverse of Eq. (34), and define $T(\lambda) = T(f(\lambda))$. Then the interior optimum satisfies

$$\frac{\mathrm{d}}{\mathrm{d}\lambda}\left[T(\lambda) + \frac{\bar{\delta}}{\lambda}\right]_{\lambda = \lambda_{\text{opt}}} = 0. \tag{38}$$

For a general $G_\theta$ this equation can be solved numerically. Below, after deriving the local stability condition, we collect the closed-form results for the two lag functions considered. In the local analysis, $\lambda_{\text{opt}}$ denotes the appropriate interior solution of Eq. (38), with the feasible boundary $f_{\text{opt}} \geq 1/2$ imposed at the end.

#### B. Cycle-growth-factor fluctuations destabilize the homogeneous optimum

The homogeneous optimum identifies the best single allocation phenotype, but it does not determine whether all cells should express that phenotype. When the cycle growth factor fluctuates, different cycles favor slightly different growth–lag compromises; a genotype that spreads its cells over nearby phenotypes can then gain from this temporal variation even if its mean remains at the homogeneous optimum.

We test this possibility by asking whether a narrow heterogeneous mutant invades the homogeneous resident. Because Eq. (34) is monotone in $f$, we use growth rate as the local phenotype coordinate and write

$$\lambda = \lambda_{\text{opt}} + \epsilon, \qquad \langle \epsilon \rangle = 0, \qquad \langle \epsilon^2 \rangle = v_\lambda. \tag{39}$$

where the brackets denote the average over phenotypes within the mutant genotype. In this coordinate, the cycle time in a realization with environmental noise $\zeta$ is

$$t(\lambda, \zeta) = T(\lambda) + \frac{\bar{\delta}(1+\zeta)}{\lambda}. \tag{40}$$

The reduced invasion term in Eq. (26) becomes

$$X(\lambda; \lambda_{\text{opt}}, \zeta) = \lambda \left[ t(\lambda_{\text{opt}}, \zeta) - t(\lambda, \zeta) \right]. \tag{41}$$

Expanding around $\lambda_{\text{opt}}$ and using the cumulant expansion in Eq. (32) gives the leading invasion fitness

$$s = \frac{v_\lambda}{2} \lambda_{\text{opt}} \left[ \lambda_{\text{opt}} \mathbb{E}_\zeta \left( t'(\lambda_{\text{opt}}, \zeta)^2 \right) - \mathbb{E}_\zeta \left( t''(\lambda_{\text{opt}}, \zeta) \right) \right] + O(v_\lambda^{3/2}). \tag{42}$$

where the first-order mean-shift term vanishes because the resident is at the homogeneous optimum. The two terms in brackets have opposite signs in their biological effect. The variance of the instantaneous selection gradient is destabilizing, because it means different phenotypes are favored in different cycles. The mean curvature term is stabilizing, because phenotypes away from the optimum pay a depletion-time cost. Thus the homogeneous optimum is locally unstable to phenotypic heterogeneity when

$$\lambda_{\text{opt}} \mathbb{E}_\zeta \left[ t'(\lambda_{\text{opt}}, \zeta)^2 \right] > \mathbb{E}_\zeta \left[ t''(\lambda_{\text{opt}}, \zeta) \right]. \tag{43}$$

The derivatives of Eq. (40) make the role of environmental variability explicit:

$$t'(\lambda, \zeta) = T'(\lambda) - \frac{\bar{\delta}(1+\zeta)}{\lambda^2}. \tag{44}$$

At the homogeneous optimum, Eq. (38) gives $T'(\lambda_{\text{opt}}) = \bar{\delta}/\lambda_{\text{opt}}^2$. Therefore

$$t'(\lambda_{\text{opt}}, \zeta) = -\frac{\bar{\delta}}{\lambda_{\text{opt}}^2} \zeta, \qquad \mathbb{E}_\zeta \left[ t'(\lambda_{\text{opt}}, \zeta)^2 \right] = \frac{\bar{\delta}^2 \sigma_\zeta^2}{\lambda_{\text{opt}}^4}. \tag{45}$$

$$\mathbb{E}_\zeta \left[ t''(\lambda_{\text{opt}}, \zeta) \right] = T''(\lambda_{\text{opt}}) + \frac{2\bar{\delta}}{\lambda_{\text{opt}}^3}. \tag{46}$$

Substitution into Eq. (43) yields the critical environmental noise

$$\sigma_\zeta^2 > \sigma_{\zeta,\text{crit}}^2 = \frac{1}{\bar{\delta}} \left[ 2 + \lambda_{\text{opt}} \frac{T''(\lambda_{\text{opt}})}{T'(\lambda_{\text{opt}})} \right]. \tag{47}$$

Here $\sigma_\zeta^2 = \text{Var}(\zeta)$ is the variance of the normalized environmental noise in Eq. (30). Figure 1 shows this local threshold for the two lag functions used below.

Collecting the closed-form results for the two lag functions, the interior homogeneous optima are

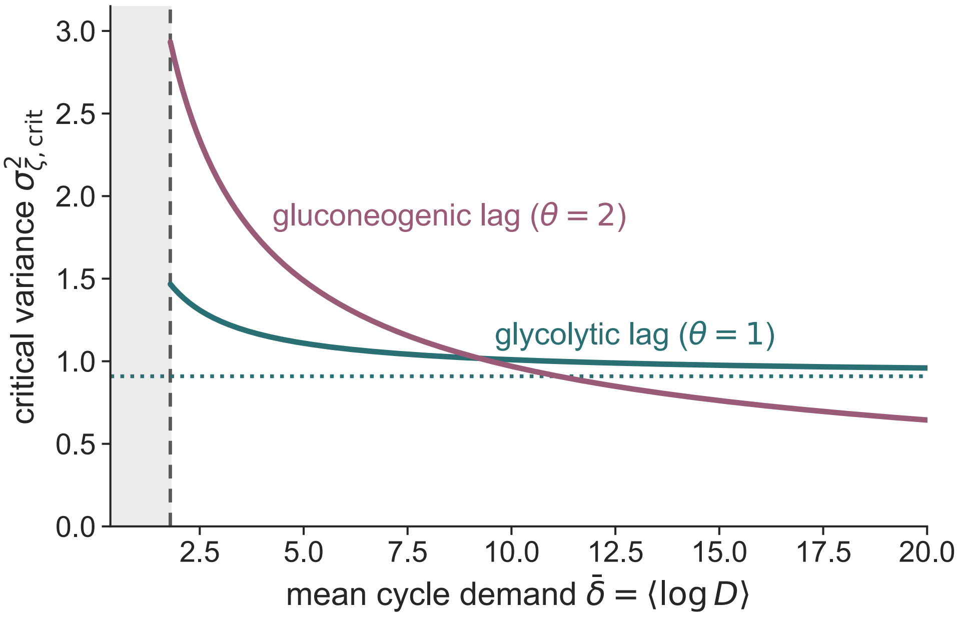


FIG. 1. **Mean cycle growth factor sets the variability threshold for phenotypic branching.** Critical growth-factor variance $(\sigma_{\zeta,\mathrm{crit}})^2$ for the glycolytic lag ($\theta = 1$, logarithmic) and the gluconeogenic lag ($\theta = 2$, inverse-preparedness), obtained from Eq. (47). Above each curve, the homogeneous optimum is locally unstable to within-genotype phenotypic variability. The shaded region marks values of $\bar{\delta}$ for which the homogeneous optimum is pinned at the boundary $f_{\mathrm{opt}} = 1/2$, so the interior branching criterion is not defined and is clipped in simulations. At large $\bar{\delta}$, the logarithmic lag retains a finite threshold, whereas the inverse-preparedness threshold vanishes.

$$\lambda^{(1)}_{\mathrm{opt}} = \frac{\lambda^\star}{1 + \tau_0 \lambda^\star / \bar{\delta}}, \qquad \lambda^{(2)}_{\mathrm{opt}} = \frac{\lambda^\star}{1 + \sqrt{\frac{\tau_0 \lambda^\star (\lambda_{\max} - \lambda^\star)}{\bar{\delta}\, \lambda_{\max}}}}, \tag{48}$$

where superscripts (1) and (2) denote $G_1(x) = -\log x$ and $G_2(x) = x^{-1} - 1$. The corresponding allocations are obtained by inverting Eq. (34),

$$f^{(\theta)}_{\mathrm{opt}} = \frac{\lambda^{(\theta)}_{\mathrm{opt}} (\lambda_{\max} - \lambda^\star)}{\lambda^\star (\lambda_{\max} - \lambda^{(\theta)}_{\mathrm{opt}})}, \qquad \theta \in \{1, 2\}, \tag{49}$$

with the boundary correction $f^{(\theta)}_{\mathrm{opt}} = 1/2$ if this interior solution falls below $1/2$. Away from that boundary, the critical variances are

$$\left(\sigma^{(1)}_{\zeta,\mathrm{crit}}\right)^2 = \frac{1}{\bar{\delta}} + \frac{1}{\lambda^\star \tau_0}, \qquad \left(\sigma^{(2)}_{\zeta,\mathrm{crit}}\right)^2 = \frac{2}{\bar{\delta}} \left[ 1 + \frac{\sqrt{\bar{\delta}}\, \lambda_{\max}}{\sqrt{(\lambda_{\max} - \lambda^\star) \lambda^\star \tau_0}} \right]. \tag{50}$$

These thresholds are local predictions for the onset of phenotypic branching around the homogeneous optimum. Within their interior domain, both critical variances decrease monotonically with the mean log growth factor $\bar{\delta}$, so increasingly severe average dilution makes phenotypic heterogeneity favorable under progressively weaker cycle-to-cycle fluctuations. The boundary at which $f_{\mathrm{opt}} = 1/2$ marks the minimum mean log growth factor for this mean-preserving local branching calculation: below this boundary, an infinitesimal phenotype distribution centered on the homogeneous optimum cannot be constructed within the feasible domain. The limits of large growth factor distinguish the two lag functions. For the inverse-preparedness lag, $\theta = 2$, $(\sigma^{(2)}_{\zeta,\mathrm{crit}})^2 \to 0$, so arbitrarily weak nonzero fluctuations eventually destabilize the homogeneous optimum. By contrast, for the logarithmic lag, $\theta = 1$, the critical variance approaches the finite value $1/(\lambda^\star \tau_0)$.

Both thresholds take a simpler form in terms of the optimal growth rate. Let $\kappa = \lambda^\star/(\lambda^\star - \lambda_{\rm opt}) = -{\rm d}\log[(1-f)/f]/{\rm d}\log\lambda$ be the elasticity of the preparedness budget $(1-f)/f$ at the optimum, and $r^{\rm opt} = (1-f_{\rm opt})/f_{\rm opt}$ the preparedness ratio of the resident. For $T = \tau_0 G_\theta[(1-f)/f]$ one finds $T' = \tau_0\kappa\,(r^{\rm opt})^{1-\theta}/\lambda$ and, at the optimum, $\lambda T''/T' = \theta\kappa - 2$. Stationarity, Eq. (38), then gives $\bar\delta = \lambda_{\rm opt}\tau_0\kappa\,(r^{\rm opt})^{1-\theta}$, and Eq. (47) becomes $\sigma^2_{\zeta,\rm crit} = \theta\kappa/\bar\delta$, that is,

$$\left(\sigma^{(1)}_{\zeta,\rm crit}\right)^2 = \frac{1}{\lambda_{\rm opt}\tau_0}, \qquad \left(\sigma^{(2)}_{\zeta,\rm crit}\right)^2 = \frac{2\,r^{\rm opt}}{\lambda_{\rm opt}\tau_0}, \tag{51}$$

which coincide with Eq. (50). For the logarithmic lag, $1/\lambda_{\rm opt} = 1/\lambda^\star + \tau_0/\bar\delta$ by Eq. (48), so that $\sigma^2_{\zeta,\rm crit} = 1/\bar\delta + 1/(\lambda^\star\tau_0)$ never falls below $\max(1/\bar\delta, 1/\lambda^\star\tau_0)$: however strong the dilution, a minimum level of environmental noise is needed.

Finite-amplitude competition and long evolutionary simulations test how this local instability appears in the nonlinear ecological dynamics. The conversion from imposed dilution and supply fluctuations to the stationary environmental-noise coordinate used in this comparison is described in Sec. V C.

### IV. MANY RESOURCES: GROWTH AND FUTURE PREPAREDNESS

With $M > 2$ resources, cells can differ both in how much they prepare for a resource switch and in which future resources they prepare for. We consider equal-quality resources with the common growth law in Eq. (11) and allow a general transition matrix. We use the allocation coordinates introduced in Eq. (3):

$$\phi_{b|b} = f_b, \qquad \phi_{a|b} = (1-f_b)q_{a|b} \quad (a \neq b), \qquad \sum_{a\neq b} q_{a|b} = 1. \tag{52}$$

The diagonal allocation $f_b$ supports growth on the present resource, while $q_{a|b}$ is the share of the remaining preparedness budget devoted to future resource $a$. The ratio

$$r_{a|b} = \frac{\phi_{a|b}}{\phi_{a|a}} = \frac{(1-f_b)q_{a|b}}{f_a}, \qquad T_{a|b} = \tau_0 G_\theta\big(\min(r_{a|b}, 1)\big) \tag{53}$$

measures how much incoming-resource machinery is already present relative to the balanced-growth target. These coordinates have distinct physiological roles: $f_a$ determines growth on the incoming resource, whereas both $f$ and $q$ determine lag through $r$. For $M = 2$, normalization fixes $q$ and leaves only the diagonal direction in the symmetric family. For $M > 2$, redistribution among futures supplies additional directions of phenotypic variation.

We test the stability of the optimal homogeneous allocation to heterogeneous mutants. The local calculations assume an interior resident and perturbations that remain on the positive-lag branch and begin growing before resident depletion. Finite distributions that encounter either physiological floor fall outside these calculations.

#### A. The homogeneous resident defines the preparedness pattern

The general invasion criterion in Eq. (27) also determines the resident mean preparedness. At fixed diagonal allocations, a change in $q_{a|b}$ changes only the lag. Differentiating the homogeneous invasion fitness and enforcing normalization within each previous-resource column gives

$$w_{a|b}\lambda_a^{\rm opt}\tau_0\left(\frac{1-f_b^{\rm opt}}{f_a^{\rm opt}}\right)^{1-\theta}\left(q_{a|b}^{\rm opt}\right)^{-\theta} = \text{constant within column } b. \tag{54}$$

Consequently,

$$q_{a|b}^{\rm opt} = \frac{\left[w_{a|b}\lambda_a^{\rm opt}(f_a^{\rm opt})^{\theta-1}\right]^{1/\theta}}{\sum_{a\neq b}\left[w_{a|b}\lambda_a^{\rm opt}(f_a^{\rm opt})^{\theta-1}\right]^{1/\theta}}, \tag{55}$$

where $\lambda_a^{\rm opt} = \lambda(f_a^{\rm opt})$ for the common growth function. This interior formula requires positive preparedness on the transition directions being varied. Within the family with one shared diagonal allocation $f$, all resident growth rates coincide and Eq. (55) reduces to

$$q_{a|b}^{\rm opt} = \frac{w_{a|b}^{1/\theta}}{\sum_{a\neq b} w_{a|b}^{1/\theta}}. \tag{56}$$

Thus logarithmic lag learns the transition probabilities conditioned on a switch, and inverse-preparedness lag learns their normalized square roots [5].

#### B. The local invasion criterion uses growth and completion time

Using the completion time already defined in Eq. (23), the one-cycle advantage is

$$X_{a|b} = \lambda_a(\boldsymbol{\phi}) \left[ t_{a|b}(\boldsymbol{\phi}_{\rm opt};\delta) - t_{a|b}(\boldsymbol{\phi};\delta) \right]. \tag{57}$$

$X_{a|b}$ is the one-cycle log-growth advantage of a phenotype relative to the homogeneous resident. A phenotype that completes the same expansion earlier has extra time to grow before resident depletion, giving $X > 0$; a slower phenotype has $X < 0$.

For a centred narrow distribution of allocation perturbations $\Delta\boldsymbol{\phi}$, the cumulant expansion of Eq. (32) gives

$$s = \frac{1}{2} \sum_b \pi_b \sum_a w_{a|b}\, \mathbb{E}_{\delta|a,b} \Big\langle \lambda_a^2 \left( \nabla t_{a|b} \cdot \Delta\boldsymbol{\phi} \right)^2 - \lambda_a\, \Delta\boldsymbol{\phi}^{\mathsf{T}} (\nabla^2 t_{a|b}) \Delta\boldsymbol{\phi} - 2 \left( \nabla \lambda_a \cdot \Delta\boldsymbol{\phi} \right) \left( \nabla t_{a|b} \cdot \Delta\boldsymbol{\phi} \right) \Big\rangle_\rho + O(\|\Delta\boldsymbol{\phi}\|^3). \tag{58}$$

In this quadratic form, $\Delta\boldsymbol{\phi}$ denotes the vector of independent allocation entries; the remaining entries are fixed by column normalization. Derivatives act on those coordinates and are evaluated at the resident. The first term rewards variation in cycle-specific completion time; the remaining terms describe the mean cost of changing completion time and growth together. For resource-dependent allocations the last term must be retained. For a shared $f$ with equal resident growth, its environmental average vanishes at homogeneous stationarity, recovering the scalar curvature–fluctuating-gradient balance of Sec. III. Self transitions are included in the sum: they have zero lag and contribute through growth alone.

Equation (58) is a quadratic form in mutant covariance. It must be non-positive for every admissible covariance for the homogeneous resident to be locally stable. Scalar thresholds therefore require a specified allocation direction. For a shared diagonal allocation the quadratic form simplifies in the variables of the next subsection, and the blocks with direct ecological meaning are then evaluated: changing investment in present growth, redistributing preparation among futures, and the coupling between the two.

#### C. The one-cycle advantage expands to second order in the growth and share deviations

From now on we assume equal-quality resources, one shared diagonal allocation $f$ per phenotype, no self transitions, and a transition-independent growth-factor distribution. Resource-dependent diagonal allocations, or growth-factor statistics that depend on the transition, require Eq. (58) itself.

Using the completion time $t = T + \delta/\lambda$ of Eq. (23), Eq. (57) reads

$$X_{a|b} = \lambda \left\{ \tau_0 \left[ G_\theta(r^{\rm opt}_{a|b}) - G_\theta(r_{a|b}) \right] + \delta \left( \frac{1}{\lambda_{\rm opt}} - \frac{1}{\lambda} \right) \right\}, \tag{59}$$

where $\lambda$ and $r_{a|b}$ are the growth rate and the preparedness ratio of the phenotype, and $\lambda_{\rm opt}$ and $r^{\rm opt}_{a|b}$ those of the resident. The first term is the lag saved with respect to the resident, the second is the difference in the time needed to grow by $\delta$, and the prefactor $\lambda$ converts time into log-growth. The phenotype enters $X$ explicitly through $\lambda$ and, through $r = (1-f)q/f$, inside the lag function $G_\theta$. We now make the dependence of $G_\theta$ on the phenotype explicit by writing it in terms of the growth rate and the shares.

The growth rate $\lambda(f)$ of Eq. (34) is monotone in $f$ and can replace it as the phenotypic coordinate, as in Sec. III: a mutant phenotype has $\lambda = \lambda_{\rm opt} + \epsilon$, with $\langle \epsilon \rangle = 0$ and $\langle \epsilon^2 \rangle = v_\lambda$ (Eq. (39)). The preparedness budget $(1-f)/f$ is then an explicit function of $\lambda$,

$$H(\lambda) \equiv \frac{1-f}{f} = \frac{\lambda_{\max}}{\lambda_{\max} - \lambda^\star} \left( \frac{\lambda^\star}{\lambda} - 1 \right), \tag{60}$$

and the preparedness ratio of Eq. (53) factorizes as $r_{a|b} = H(\lambda)\, q_{a|b}$. A phenotype is thus specified by its growth rate, which fixes growth on the incoming resource and the total preparedness budget, and by the shares $q_{a|b}$, which divide that budget among the futures.

We write $q_{a|b} = q^{\rm opt}_{a|b}(1+z_{a|b})$, with $\langle z_{a|b}\rangle = 0$ and, by normalization, $\sum_a q^{\rm opt}_{a|b} z_{a|b} = 0$ for every $b$. Expanding Eq. (59) to second order around the resident, with $r_{a|b} = H(\lambda)\, q_{a|b}$ and $G_\theta(r) = (r^{1-\theta}-1)/(\theta-1)$, gives $X_{a|b} = X^{[1]}_{a|b}+X^{[2]}_{a|b}+O(3)$. Explicit calculations lead to

$$X^{[1]}_{a|b} = \lambda_{\rm opt}\tau_0 \left(r^{\rm opt}_{a|b}\right)^{1-\theta} z_{a|b} + \left[\frac{\delta}{\lambda_{\rm opt}} - \tau_0 \left(r^{\rm opt}_{a|b}\right)^{1-\theta} \frac{\lambda^\star}{\lambda^\star - \lambda_{\rm opt}}\right]\epsilon, \tag{61}$$

$$X^{[2]}_{a|b} = -\tfrac{1}{2}\theta\tau_0 \left(r^{\rm opt}_{a|b}\right)^{1-\theta} \left[\lambda_{\rm opt} z^2_{a|b} + \left(\frac{\lambda^\star}{\lambda^\star - \lambda_{\rm opt}}\right)^2 \frac{\epsilon^2}{\lambda_{\rm opt}}\right] + \tau_0 \left(r^{\rm opt}_{a|b}\right)^{1-\theta} \left[1 + (\theta-1)\frac{\lambda^\star}{\lambda^\star - \lambda_{\rm opt}}\right] \epsilon\, z_{a|b}. \tag{62}$$

The growth-rate deviation $\epsilon$ is a property of the phenotype and is the same on every transition, whereas $z_{a|b}$ and $r^{\rm opt}_{a|b}$ refer to the transition realized in the cycle: no sum over targets appears in $X$, and the average over transitions is taken in Eq. (65). The factor $(r^{\rm opt})^{1-\theta}$ is the lag cost, in units of $\tau_0$, of a relative loss of preparedness, $-\tau_0^{-1}\partial T/\partial \log r$ at the resident, and $\kappa = \lambda^\star/(\lambda^\star - \lambda_{\rm opt}) = -d\log H/d\log\lambda$ is the elasticity of the preparedness budget at the resident. The $z^2$ and $\epsilon^2$ terms of $X^{[2]}$ are the curvature of the lag law in $\log r$, and the cross term comes from the growth-rate prefactor of Eq. (59) multiplying the first-order change of $r$, together with the second-order change of the budget. Because the growth rate is itself the coordinate, no derivative of the growth law appears.

For the glycolytic lag, $\theta = 1$ and $G_{\rm glyc}(r) = -\log r$, the resident ratio drops out:

$$X^{[1]}_{\rm glyc} = \lambda_{\rm opt}\tau_0\, z + \left[\frac{\delta}{\lambda_{\rm opt}} - \tau_0\kappa\right]\epsilon,$$

$$X^{[2]}_{\rm glyc} = -\tfrac{1}{2}\tau_0 \left[\lambda_{\rm opt} z^2 + \kappa^2 \frac{\epsilon^2}{\lambda_{\rm opt}}\right] + \tau_0\, \epsilon\, z. \tag{63}$$

For the gluconeogenic lag, $\theta = 2$ and $G_{\rm gneo}(r) = 1/r - 1$,

$$X^{[1]}_{\rm gneo} = \frac{\lambda_{\rm opt}\tau_0}{r^{\rm opt}}\, z + \left[\frac{\delta}{\lambda_{\rm opt}} - \frac{\tau_0}{r^{\rm opt}}\kappa\right]\epsilon,$$

$$X^{[2]}_{\rm gneo} = -\frac{\tau_0}{r^{\rm opt}} \left[\lambda_{\rm opt} z^2 + \kappa^2 \frac{\epsilon^2}{\lambda_{\rm opt}}\right] + \frac{\tau_0}{r^{\rm opt}}\left[1+\kappa\right] \epsilon\, z, \tag{64}$$

with the indices $a|b$ of $X$, $z$ and $r^{\rm opt}$ left implicit. The gluconeogenic expressions follow from the glycolytic ones by replacing $\tau_0$ with $\tau_0/r^{\rm opt}$ in every term, doubling the curvature terms, and multiplying the cross term by $1+\kappa$.

#### D. The glycolytic invasion fitness is the sum of a preparedness term and a growth term

The perturbations are centred, so $\langle X^{[1]}\rangle = 0$ and Eq. (58) reduces to

$$s = \sum_b \pi_b \sum_{a\neq b} w_{a|b}\, \mathbb{E}_\delta \left[\left\langle X^{[2]}\right\rangle + \tfrac{1}{2}\left\langle \left(X^{[1]}\right)^2\right\rangle\right] + O(3). \tag{65}$$

We evaluate it with $\langle \epsilon\rangle = \langle z_{a|b}\rangle = 0$, $\langle \epsilon^2\rangle = v_\lambda$ and $\langle z^2_{a|b}\rangle = v_{z,a|b}$, which is ${\rm Var}_\rho(q_{a|b})/(q^{\rm opt}_{a|b})^2$ and depends on the transition through the target. In full generality, the covariance $\langle \epsilon\, z_{a|b}\rangle$ is kept as well. The growth factor has mean $\bar\delta$ and variance $\bar\delta^2\sigma^2_\zeta$, and is independent of the phenotype and of the transition.

One further simplification comes from the optimality of the resident in the growth rate. The resident minimizes the mean completion time, $\mathbb{E}[\partial t/\partial\lambda] = 0$. Since $X = \lambda(t_{\rm opt} - t)$, the coefficient of $\epsilon$ in $X^{[1]}$ is $-\lambda_{\rm opt}\, \partial t/\partial\lambda$, so its average over the growth factor and transitions vanishes, implying that the mean log growth factor $\bar\delta$ satisfies

$$\bar\delta = \lambda_{\rm opt}\tau_0\, \kappa \sum_b \pi_b \sum_{a\neq b} w_{a|b} \left(r^{\rm opt}_{a|b}\right)^{1-\theta}. \tag{66}$$

For the glycolytic lag the sum equals one, so $\bar\delta = \lambda_{\rm opt}\tau_0\kappa$ on every transition, which is equivalent to $1/\lambda_{\rm opt} = 1/\lambda^\star + \tau_0/\bar\delta$. In this case the coefficient of $\epsilon$ in $X^{[1]}$ then reduces to $(\delta - \bar\delta)/\lambda_{\rm opt}$, which has zero mean and variance $\bar\delta^2\sigma^2_\zeta/\lambda^2_{\rm opt}$.

For the glycolytic lag, Eq. (63) gives

$$\left\langle X^{[2]}_{\rm glyc}\right\rangle = -\tfrac{1}{2}\tau_0\lambda_{\rm opt}\, v_{z,a|b} - \frac{\tau_0\kappa^2}{2\lambda_{\rm opt}}\, v_\lambda + \tau_0 \left\langle \epsilon\, z_{a|b}\right\rangle ,$$

$$\tfrac{1}{2}\mathbb{E}\left\langle \left(X^{[1]}_{\rm glyc}\right)^2\right\rangle = \tfrac{1}{2}\lambda_{\rm opt}^2\tau_0^2\, v_{z,a|b} + \frac{\bar\delta^2\sigma_\zeta^2}{2\lambda_{\rm opt}^2}\, v_\lambda .$$

The covariance with each single target need not vanish. However, the covariance term vanishes after the sum over targets, as a consequence of the normalization of the preparedness weights: a cell that prepares more for one future must prepare less for the others, and since $q^{\rm opt}_{a|b} = w_{a|b}$ the weighted deviations $w_{a|b}z_{a|b} = q_{a|b} - q^{\rm opt}_{a|b}$ add up to zero for every phenotype, whatever its correlation with $\epsilon$,

$$\tau_0 \sum_b \pi_b \sum_{a\neq b} w_{a|b} \left\langle \epsilon\, z_{a|b}\right\rangle = \tau_0 \sum_b \pi_b \left\langle \epsilon \sum_{a\neq b} \left(q_{a|b} - q^{\rm opt}_{a|b}\right)\right\rangle = 0. \tag{67}$$

Hence

$$s_{\rm glyc} = \tfrac{1}{2}\lambda_{\rm opt}\tau_0\left(\lambda_{\rm opt}\tau_0 - 1\right)\sum_b \pi_b \sum_{a\neq b} w_{a|b}\, v_{z,a|b} + \frac{\bar\delta^2}{2\lambda_{\rm opt}^2}\left(\sigma_\zeta^2 - \frac{1}{\lambda_{\rm opt}\tau_0}\right) v_\lambda . \tag{68}$$

The sum over targets is the mean $\chi^2$ distance of the cells' preparedness from the resident. The individual $v_{z,a|b}$ depend on the target, but for a Dirichlet distribution of the weights around $q^{\rm opt}$ with concentration $1/\sigma_q^2$ the factor $1/q^{\rm opt}$ in $v_z$ cancels the weight, and for any transition matrix

$$s_{\rm glyc} = \tfrac{1}{2}\lambda_{\rm opt}\tau_0\left(\lambda_{\rm opt}\tau_0 - 1\right)(M-2)\frac{\sigma_q^2}{1+\sigma_q^2} + \frac{\bar\delta^2}{2\lambda_{\rm opt}^2}\left(\sigma_\zeta^2 - \frac{1}{\lambda_{\rm opt}\tau_0}\right) v_\lambda . \tag{69}$$

Thus, the local stability of the homogeneous resident is determined by two additive contributions: the first term is the effect of redistributing preparedness among future resources, and the second is the effect of changing growth on the present resource.

Interestingly, two thresholds appear, one for each block of the quadratic form, and they are both set by the same number, the lag time in units of the growth time, $\lambda_{\rm opt}\tau_0$.

The preparedness term does not depend on the growth factor and is positive if and only if $\lambda_{\rm opt}\tau_0 > 1$: all-in hedging pays when the lag, measured in growth times, exceeds one. At $f = f_{\rm opt}$, the smooth factor by which a cell with weight $q^{\rm opt}(1+z)$ on the incoming resource multiplies its biomass relative to the resident is $e^X = (1+z)^{\lambda_{\rm opt}\tau_0}$, convex in the weights if $\lambda_{\rm opt}\tau_0 > 1$ and concave if $\lambda_{\rm opt}\tau_0 < 1$. For $\lambda_{\rm opt}\tau_0 > 1$ its average at fixed mean weights is therefore largest when the weights sit on the vertices of the simplex, that is, when each cell prepares for a single future.

The growth term is the only place where $\sigma_\zeta$ enters. It is positive if and only if $\sigma_\zeta^2\lambda_{\rm opt}\tau_0 > 1$, that is, above

$$\sigma^2_{\zeta,\rm crit} = \frac{1}{\lambda_{\rm opt}\tau_0}. \tag{70}$$

By stationarity this equals $1/\bar\delta + 1/(\lambda^\star\tau_0)$, the threshold of the two-resource problem. It also shows that $\lambda_{\rm opt}\tau_0$ is smaller than both $\bar\delta$ and $\lambda^\star\tau_0$, so that $\lambda_{\rm opt}\tau_0 > 1$ requires strong dilution and a lag longer than the growth time at the same time.

Two regimes follow. For $\lambda_{\rm opt}\tau_0 < 1$ only the growth term can be positive, and only if $\sigma_\zeta > (\lambda_{\rm opt}\tau_0)^{-1/2} > 1$, that is, if the growth factor fluctuates by more than its mean. For $\lambda_{\rm opt}\tau_0 > 1$ the $z$ term is positive at any noise level, and the growth term as soon as $\sigma_\zeta > (\lambda_{\rm opt}\tau_0)^{-1/2}$, which is below one and goes to zero as $\lambda_{\rm opt}\tau_0$ grows: at moderate noise both terms are positive. In the plane $(\lambda_{\rm opt}\tau_0, \sigma_\zeta)$ (Fig. 2, left) the $z$ term is positive to the right of $\lambda_{\rm opt}\tau_0 = 1$ and the growth term above $\sigma_\zeta = (\lambda_{\rm opt}\tau_0)^{-1/2}$. The two boundaries meet at $(1,1)$ and delimit four regions: neither term, only the growth term, only the $z$ term, and both.

#### E. The gluconeogenic lag keeps the same two terms but its cost depends on the target

The calculation follows that of the glycolytic lag: Eq. (65) is evaluated with the expansion of Eq. (64), the growth factor is averaged, and the terms in $z$ and in $\epsilon$ are collected. Two things change. First, a relative loss of preparedness costs a lag $\tau_0/r^{\rm opt}_{a|b}$ instead of $\tau_0$, so the cost depends on the target through the resident preparedness, and the curvature of the lag in $\log r$ doubles. Second, the sum in Eq. (66) is no longer one but the average of $1/r^{\rm opt}_{a|b}$ over transitions, so stationarity holds only on average: the growth coefficient in $X^{[1]}$ has zero mean over transitions but not on each of them, and its fluctuation between transitions adds to that of the growth factor.

Collecting the terms of Eq. (65), with stationarity holding only on average, gives

$$s_{\rm gneo} = \mathbb{E}_{b,a}\left\{\tfrac{1}{2}\frac{\lambda_{\rm opt}\tau_0}{r^{\rm opt}}\left(\frac{\lambda_{\rm opt}\tau_0}{r^{\rm opt}}-2\right)v_{z,a|b} + \frac{\tau_0}{r^{\rm opt}}\left[1+\kappa+\bar\delta-\frac{\lambda_{\rm opt}\tau_0\kappa}{r^{\rm opt}}\right]\langle \epsilon\, z_{a|b}\rangle\right\} + \frac{\bar\delta^2}{2\lambda_{\rm opt}^2}\left\{\sigma_\zeta^2+\frac{{\rm Var}_{b,a}(1/r^{\rm opt})}{\mathbb{E}_{b,a}[1/r^{\rm opt}]^2}-\frac{2}{\lambda_{\rm opt}\tau_0\,\mathbb{E}_{b,a}[1/r^{\rm opt}]}\right\}v_\lambda, \tag{71}$$

where $\mathbb{E}_{b,a}$ is the average over transitions with weights $\pi_b w_{a|b}$ and $r^{\rm opt}=r^{\rm opt}_{a|b}$ inside it. With respect to Eq. (68) the coefficient of the preparedness term depends on the target, the growth term acquires the variance of $1/r^{\rm opt}$ over transitions, which acts as additional noise in the coefficient of $\epsilon$, and the coefficient of the covariance depends on the target, also through the deviation of each transition from stationarity.

In a symmetric environment (Sec. IV F) all transitions have the same $r^{\rm opt}$, the variance vanishes, and stationarity holds on every transition. The covariance has the same coefficient on every target and, since $q^{\rm opt}_{a|b}=w_{a|b}$, it vanishes after the sum over targets, as in Eq. (67). Thus, Eq. (71) reduces to

$$s_{\rm gneo} = \tfrac{1}{2}\frac{\lambda_{\rm opt}\tau_0}{r^{\rm opt}}\left(\frac{\lambda_{\rm opt}\tau_0}{r^{\rm opt}}-2\right)\sum_b \pi_b\sum_{a\neq b} w_{a|b}\,v_{z,a|b} + \frac{\bar\delta^2}{2\lambda_{\rm opt}^2}\left(\sigma_\zeta^2-\frac{2r^{\rm opt}}{\lambda_{\rm opt}\tau_0}\right)v_\lambda. \tag{72}$$

This has the structure of Eq. (68), with $\lambda_{\rm opt}\tau_0$ replaced by $\lambda_{\rm opt}\tau_0/r^{\rm opt}$ and the number 1 in both brackets replaced by 2. For a Dirichlet distribution of the weights around $q^{\rm opt}$ with concentration $1/\sigma_q^2$ the sum over targets is the same mean $\chi^2$ distance as in the glycolytic case, and

$$s_{\rm gneo} = \tfrac{1}{2}\frac{\lambda_{\rm opt}\tau_0}{r^{\rm opt}}\left(\frac{\lambda_{\rm opt}\tau_0}{r^{\rm opt}}-2\right)(M-2)\frac{\sigma_q^2}{1+\sigma_q^2} + \frac{\bar\delta^2}{2\lambda_{\rm opt}^2}\left(\sigma_\zeta^2-\frac{2r^{\rm opt}}{\lambda_{\rm opt}\tau_0}\right)v_\lambda. \tag{73}$$

The preparedness term is again independent of the growth factor.

In the symmetric environment, and only there, the gluconeogenic lag has, as the glycolytic lag, two thresholds set by the same number, now $\lambda_{\rm opt}\tau_0/r^{\rm opt}$. At fixed $\lambda_{\rm opt}\tau_0$ both thresholds are those of the glycolytic lag multiplied by $2r^{\rm opt}$, so the gluconeogenic lag is more permissive if $r^{\rm opt}<\frac{1}{2}$.

Finally, for structured transitions the variance of $1/r^{\rm opt}$ between transitions lowers the critical noise of the growth term to $\sigma^2_{\zeta,\rm crit}=2\kappa/\bar\delta-{\rm Var}_{b,a}(1/r^{\rm opt})/\mathbb{E}_{b,a}[1/r^{\rm opt}]^2$, while the covariance no longer cancels and mixed directions require the full quadratic form of Eq. (58).

A closed form remains for circulant transitions, in which the switch from $b$ goes to $b+k$ (modulo $M$) with probability $w_k$ independent of $b$, so that the stationary distribution is uniform and, at the optimum, the weights are $q^{\rm opt}_k=\sqrt{w_k}/S$ with $S=\sum_k\sqrt{w_k}$ and $r^{\rm opt}_k=(1-f)q^{\rm opt}_k/f$. The mean lag, $\tau_0[\,fS^2/(1-f)-1]$, depends on the transitions only through $S^2$, which replaces $M-1$ in the symmetric case; the threshold of the growth term is thus $\sigma_\zeta^2\,\lambda_{\rm opt}\tau_0/r^{\rm opt}=2$ for every $w_k$, with $r^{\rm opt}=(1-f)/(S^2f)$. The preparedness term couples only the weights of the target of the switch: a mutant whose weight on the target $k$ is $q^{\rm opt}_k(1+z)$ completes the cycle earlier by $\tau_0(1-1/(1+z))/r^{\rm opt}_k$, and Eq. (71) gives for each target the contribution $\frac{1}{2}(\lambda_{\rm opt}\tau_0/r^{\rm opt}_k)(\lambda_{\rm opt}\tau_0/r^{\rm opt}_k-2)\,v_{z,k}$, where $v_{z,k}\propto(1-q^{\rm opt}_k)/q^{\rm opt}_k$ for a Dirichlet distribution. Summing over the targets with weights $w_k$, the preparedness term changes sign at $x=x^*$, with $x=\lambda_{\rm opt}\tau_0/r^{\rm opt}$ and

$$x^* = \frac{2(M-2)S^2}{S\sum_k w_k^{-1/2}-(M-1)}, \tag{74}$$

which is 2 for uniform $w_k$ and decreases as the transitions become predictable (Fig. 4). It holds as long as the lag is not clipped at zero for any target, $\max_k r^{\rm opt}_k<1$.

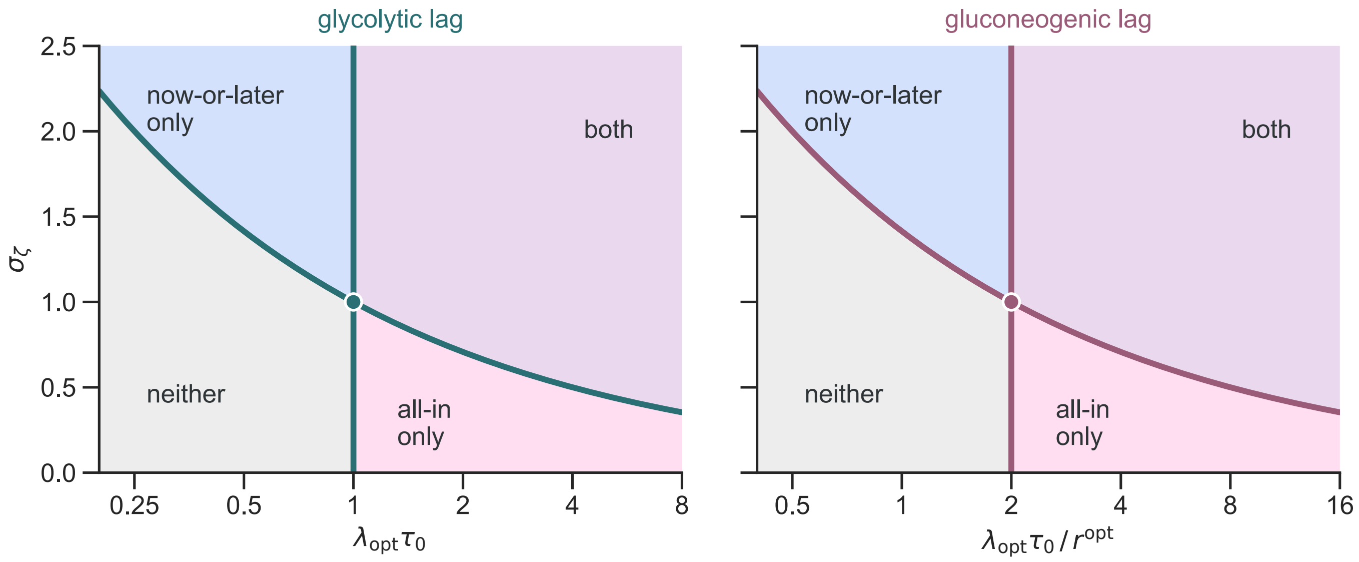


FIG. 2. **A single number, the lag time in units of the growth time, organizes the local stability of the homogeneous resident for both strategies.** Left, glycolytic lag: the preparedness term (all-in hedging) is positive to the right of $\lambda_{\rm opt}\tau_0 = 1$, and the growth term (now-or-later hedging) above $\sigma_\zeta = (\lambda_{\rm opt}\tau_0)^{-1/2}$. The two boundaries meet at $(1, 1)$ and delimit four regions; for $\lambda_{\rm opt}\tau_0 > 1$ the threshold of the growth term is below one, so that both terms are positive at moderate noise. Right, gluconeogenic lag: the same map in terms of $\lambda_{\rm opt}\tau_0/r^{\rm opt}$, with the threshold of the preparedness term at 2 instead of 1; the boundaries still meet at $\sigma_\zeta = 1$. Smooth branch, equal-quality resources, symmetric transitions for the gluconeogenic lag; boundaries are analytic, Eqs. (68) and (72).

#### F. For symmetric resources the optimum and the thresholds take closed forms

Assume equal-quality resources, no self transitions, uniform switches, and a stationary growth-factor distribution independent of the resource:

$$w_{a|b} = \frac{1}{M-1} \quad (a \neq b), \qquad \phi_{b|b} = f, \qquad \phi_{a|b} = \frac{1-f}{M-1}. \tag{75}$$

With the preparedness weights held at $q^{\rm opt} = 1/(M-1)$, $f$ is the only coordinate: growth is Eq. (34) and lag is

$$T_M^{(\theta)}(f) = \tau_0 G_\theta\!\left(\frac{1-f}{(M-1)f}\right). \tag{76}$$

The optimum follows as in Sec. III, and the growth rate and preparedness ratio of the resident are

$$\lambda_{\rm opt}^{\rm glyc} = \frac{\lambda^\star}{1+\tau_0\lambda^\star/\bar{\delta}}, \qquad r_{\rm glyc}^{\rm opt} = \frac{\lambda_{\max}\lambda^\star\tau_0}{(M-1)\bar{\delta}(\lambda_{\max}-\lambda^\star)},$$
$$\lambda_{\rm opt}^{\rm gneo} = \frac{\lambda^\star}{1+\sqrt{(M-1)\tau_0\lambda^\star(\lambda_{\max}-\lambda^\star)/(\bar{\delta}\lambda_{\max})}}, \qquad r_{\rm gneo}^{\rm opt} = \sqrt{r_{\rm glyc}^{\rm opt}}. \tag{77}$$

The smooth branch requires $r^{\rm opt} < 1$, that is, $f_{\rm opt} > 1/M$, which for both lags means

$$\bar{\delta} > \bar{\delta}_{{\rm b},M} = \frac{\tau_0\lambda_{\max}\lambda^\star}{(M-1)(\lambda_{\max}-\lambda^\star)}. \tag{78}$$

Below it $f_{\rm opt} = 1/M$, where all resource programs receive equal allocation and lag is zero.

In the symmetric environment, Eqs. (68) and (72) give the critical noise of the growth term

$$\left(\sigma_{\zeta,{\rm crit},M}^{\rm glyc}\right)^2 = \frac{1}{\bar{\delta}} + \frac{1}{\lambda^\star\tau_0} = \frac{1}{\lambda_{\rm opt}^{\rm glyc}\tau_0}, \tag{79}$$

$$\left(\sigma_{\zeta,{\rm crit},M}^{\rm gneo}\right)^2 = \frac{2}{\bar{\delta}}\left[1+\frac{\sqrt{\bar{\delta}}\,\lambda_{\max}}{\sqrt{(M-1)(\lambda_{\max}-\lambda^\star)\lambda^\star\tau_0}}\right] = \frac{2\,r_{\rm gneo}^{\rm opt}}{\lambda_{\rm opt}^{\rm gneo}\tau_0}. \tag{80}$$

In the interior, the glycolytic threshold does not depend on $M$, because dividing preparation among $M-1$ alternatives only adds a constant to the logarithmic lag, whereas the gluconeogenic threshold decreases with $M$, because it multiplies the allocation-dependent part of the inverse-preparedness lag by $M-1$.

### V. NUMERICAL SIMULATIONS

#### A. Simulation Methods

The simulations implement the growth–dilution map of Sec. I at the level of explicit ecological cycles. Each run generates a realized environmental history, expresses each genotype as a finite ensemble of phenotypes, solves resource depletion in that phenotype population, aggregates biomass back to genotypes, and then applies the prescribed invasion or mutation protocol. The same ecological simulator is used for fixed-strategy competitions and for evolutionary simulations; only the allowed mutation axes and the scoring rule change between protocols.

Each cycle begins immediately after dilution from the previous cycle. The next resource identity is drawn from the transition matrix in Eq. (6), and the cycle draws a dilution factor $D^d$ and a resource supply $c_0^d$. In the baseline two-resource simulations, $c_0^d$ is fixed and $\log D^d$ fluctuates. In the random-supply extensions, both variables may fluctuate. The environmental draw sets the realized transition $b \to a$ and the amount of biomass that must be produced before the next dilution.

For each expressed phenotype, the simulator computes the growth rate on the supplied resource and the lag after the environmental switch from the same allocation-dependent growth–lag functions used in the analytic model. The ecological solver then finds the depletion time $t_s^d$ at which the total biomass produced by all phenotype packets equals the supplied resource. Phenotypes whose lag exceeds $t_s^d$ do not grow in that cycle; phenotypes that start before depletion expand exponentially for the remaining time. After depletion, all phenotype biomasses are divided by $D^d$.

After growth and dilution, all surviving phenotype packets are aggregated back to their parent genotype. This means that selection does not act on the success of any individual phenotype, but rather on the total biomass of the genotype, which is the sum of the biomasses of all its expressed phenotypes. A heterogeneous genotype therefore succeeds only if the combined biomass of the phenotypes it expresses increases relative to competitors.

To reduce the computational cost of simulating heterogeneous genotypes, the simulator represents each genotype as a finite ensemble of phenotype packets. In particular, for a genotype with a mean allocation matrix $\bar{\boldsymbol{\phi}}_g$ and variability parameters $\sigma_f$ and $\sigma_q$, the simulator samples a finite number of phenotype packets from the corresponding distribution of allocations. Each packet represents a subpopulation of cells that share the same allocation strategy, and the total biomass of the genotype is divided equally among these packets before growth. After growth and dilution, the biomass of each packet is aggregated back to the parent genotype, and the process repeats for the next cycle. The $N_{\text{pheno}}$ packets of a genotype are drawn afresh and independently at the beginning of every cycle, so phenotypes carry no memory from one cycle to the next, and a genotype with $\sigma_f = \sigma_q = 0$ is represented by a single packet.

A genotype is therefore heritable both at the level of a mean allocation matrix and the variability parameters $\sigma_f$, which controls the variability in the diagonal allocation to the currently supplied resource, and $\sigma_q$, which controls the variability in the off-diagonal allocation to other resources. Naturally, the latter is only relevant for $M > 2$, so that phenotypes with the same current-resource allocation differ in how they distribute preparedness across possible future resources.

For two resources, the sampled coordinate is the current-resource allocation $f$, drawn from a beta distribution with mean $\bar{f}$ and variance $\sigma_f^2$, that is, with shape parameters

$$\alpha_f = \bar{f}\chi, \qquad \beta_f = (1-\bar{f})\chi, \qquad \chi = \frac{\bar{f}(1-\bar{f})}{\sigma_f^2} - 1, \tag{81}$$

which requires $\sigma_f < \sigma_f^{\max} = \sqrt{\bar{f}(1-\bar{f})}$. For $M > 2$ the same $f$ is shared by all columns of a phenotype.

For $M > 2$, off-diagonal variability is implemented in the normalized preparedness weights $q_{a|b}$ of Eq. (3). After drawing the diagonal allocation $f$ for a phenotype, the simulator treats each previous-resource column $b$ independently and draws the vector $q_{\cdot|b}$ over $a \neq b$ from a Dirichlet distribution with mean equal to the genotype mean $\bar{q}_{a|b}$:

$$q_{\cdot|b} \sim \text{Dirichlet}\left(\alpha_{a|b}\right), \qquad \alpha_{a|b} = \frac{\bar{q}_{a|b}}{\sigma_q^2} \quad (a \neq b). \tag{82}$$

Thus $\sigma_q$ is a Dirichlet dispersion parameter for the off-diagonal allocation fractions around a fixed mean. The total Dirichlet concentration is $\alpha_0 = 1/\sigma_q^2$, and each off-diagonal component has variance

$$\text{Var}(q_{a|b}) = \frac{\bar{q}_{a|b}(1-\bar{q}_{a|b})}{\alpha_0 + 1} = \frac{\bar{q}_{a|b}(1-\bar{q}_{a|b})\sigma_q^2}{1+\sigma_q^2}. \tag{83}$$

The expressed allocation is reconstructed as $\phi_{a|b} = (1-f)q_{a|b}$ for $a \neq b$ and $\phi_{b|b} = f$. Population summaries use the biomass-weighted genotype-level dispersions

$$\bar{\sigma}_f^{B,d} = \frac{\sum_g B_g^d \sigma_{f,g}}{\sum_g B_g^d}, \tag{84}$$

$$\bar{\sigma}_q^{B,d} = \frac{\sum_g B_g^d \sigma_{q,g}}{\sum_g B_g^d}, \tag{85}$$

where $B_g^d$ is the biomass of genotype $g$ at cycle $d$.

Feasibility is enforced before phenotypes are sampled. The requested variability in $f$ must be compatible with a distribution supported on $0 \le f \le 1$. When a mutation would place the mean too close to the boundary for the requested variability, the mean allocation is projected back into the feasible region while preserving the off-diagonal proportions.

#### B. Fixed-strategy assays and evolutionary runs use the same simulator and differ only in the protocol

In fixed-strategy assays, prescribed genotypes start at prescribed biomass fractions, with equal biomass in direct competitions and the mutant at relative abundance $10^{-7}$ in rare-mutant assays, and neither mutation nor pruning occurs. Genotypes compared at a given noise and replicate share the same environmental history, which is drawn from an environmental seed common to all genotypes, whereas the phenotype sampling of each run has its own seed, so that responses to the traits are not confounded with different environmental samples. The outcome is scored from the log-ratio of the genotype biomasses,

$$\ell^d = \log \frac{B_{\text{het}}^d + \epsilon}{B_{\text{hom}}^d + \epsilon}, \tag{86}$$

with $\epsilon$ a small floor that keeps the logarithm finite, or, in rare-mutant assays, from its change relative to the initial state,

$$\Delta\ell^d = \log \frac{B_{\text{mut}}^d + \epsilon}{B_{\text{res}}^d + \epsilon} - \log \frac{B_{\text{mut}}^0 + \epsilon}{B_{\text{res}}^0 + \epsilon}. \tag{87}$$

A run stops early when this log-ratio stays beyond a fixed threshold over a prescribed window after a minimum number of cycles, and otherwise at a maximum number of cycles, where it is classified only if its final tail lies beyond the threshold and is left unresolved if not. The win probability is the fraction of decided histories won by a genotype.

In evolutionary runs, mutants are generated sequentially. In each cycle, with probability $1/\tau_{\text{mut}}$ (the mutation interval $\tau_{\text{mut}}$ is the mean number of cycles between mutation events), a parent is drawn with probability proportional to its positive offspring production in that cycle, the biomass produced before dilution minus the biomass inherited at its start, so that lineages that persist through dilution without producing net offspring do not seed mutants. The mutant is created from the parent by the kernels below and enters at relative abundance $10^{-7}$. After growth, dilution and the aggregation of phenotypes, genotypes with relative abundance below the pruning threshold, $10^{-12}$, are removed, and at most $K$ lineages, the most abundant ones, are retained. The mutation kernels act on latent coordinates that keep every genotype feasible. The mean diagonal allocation $f$ is mutated through its logit with a Gaussian step $\sigma_{m,\phi}$. The variability $\sigma_f$ is mutated through a nonnegative latent amplitude $h$, with $\sigma_f = \sigma_f^{\max}(f)\, h/\sqrt{1+h^2}$ and a Gaussian step $\sigma_{m,\sigma_f}$ reflected at zero, so that $\sigma_f = 0$ is not an absorbing state. The dispersion $\sigma_q$ is mutated by a log-normal step $\sigma_{m,\sigma_q}$, with values below $10^{-6}$ set to zero and an upper bound $\sigma_q \le 2$, and the mean off-diagonal weights by a multiplicative log-normal step renormalized within each column. In the independent protocol every event mutates the mean allocation, $\sigma_f$ and $\sigma_q$; in the one-axis protocol an event mutates either the mean allocation, with probability $p_f$, or $\sigma_f$, but not both.

Population summaries are biomass-weighted averages over the surviving genotypes, $\bar{f}^B$, $\bar{\sigma}_f^B$ (Eq. (84)) and $\bar{\sigma}_q^B$ (Eq. (85)), together with the values of the dominant lineage. For preparedness we also report the effective number of prepared futures $e^H$, with $H = -\sum_{a\neq b} q_{a|b} \log q_{a|b}$ the entropy of the off-diagonal weights of a phenotype, averaged over phenotypes reconstructed from the biomass-weighted endpoint genotypes. Tail values are averages over the final analysis window of each run.

Several choices of the simulator are numerical controls rather than parameters of the model: the number of phenotype packets $N_{\text{pheno}}$, the mutation steps and interval, the seeding abundance, the pruning threshold, the lineage cap $K$, the upper bound on $\sigma_q$, the maximum number of cycles and the tail window. They are varied wherever a conclusion depends on them, and Table I lists their values in the simulations of the main text. Two effects deserve explicit mention. First, a finite phenotype sample gives a rare mutant a bonus: a genotype identical to the resident, whose own sample is not tuned to the depletion time fixed by the resident, gains a positive growth difference per cycle that scales approximately as $1/N_{\text{pheno}}$ and grows with $M$; in a many-resource test at $\sigma_D = 2.4$ and $\tau_0 = 3$ with $N_{\text{pheno}} = 101$ it was 0.006 per cycle for $M = 5$ and 0.015 for $M = 10$. It matters only when the effect measured is comparably small, as near thresholds, and is expected in evolutionary runs, where every new genotype draws independent phenotypes. Second, the upper bound on $\sigma_q$ limits how close cells can come to being specialists, and it is the control that most affects the endpoints of the many-resource evolutionary runs of the main text.

TABLE I. **Settings of the simulations of Figs. 2–5 of the main text.** Common to all: $\lambda^\star = 1.1$, $\lambda_{\max} = 2.85$, random dilution with fixed supply; glycolytic lag with $\tau_0 = 1$ unless stated ($\lambda_{\text{opt}} = 0.55$ in 4C, $\tau_0$ set by $\lambda_{\text{opt}}\tau_0$ at $\bar{\delta} = 3.5$ in 4E and 5, gluconeogenic also in 4D); uniform switches without self transitions for $M > 2$. In 4E the noise is $\sigma_\zeta = 0.4$ ($\sigma_D = 0.416$) and the runs start from the homogeneous optimum with $\sigma_f = 10^{-3}$ and $\sigma_q = 0.1$; in 5, $\sigma_\zeta = 0.4$ for the neither and all-in regions and 1.6 ($\sigma_D = 1.67$) for the now-or-later and mixed regions, with $(\sigma_f, \sigma_q) = (0.01, 0.1)$, $(0.15, 0.1)$, $(0.01, 3)$ and $(0.15, 3)$ and the same mean allocation. Histories are per grid point and tail is the scoring window. Runs with $\leq$ are adaptive: they stop when the log-ratio stays beyond $\pm 6$ over 750 cycles (1500 in 4D) after $3 \times 10^3$ ($10^4$) cycles; 4C stops when one genotype holds more than 0.999 of the biomass for 300 cycles after $10^3$.

| Fig. | assay | $M$ | $\bar{\delta}$ | $N_{\text{pheno}}$ | histories | cycles | mutation |
|---|---|---|---|---|---|---|---|
| 2A,B | competition | 2 | 3 | 201 | 50–200 | $\leq 2 \times 10^4$, tail $2.5 \times 10^3$ | none |
| 2C | rare mutant | 2 | $\log 7$ | 101 | 10–40 | $\leq 2 \times 10^4$, tail $2.5 \times 10^3$ | none |
| 3 | evolution, one axis | 2 | $\log 7$ | 201 | 5 | $4 \times 10^6$, tail $8 \times 10^5$ | $p_f = \frac{1}{4}$, $\sigma_{m,\phi} = 0.01$, $\sigma_{m,\sigma_f} = 0.0025$, $\tau_{\text{mut}} = 10$, $K = 50$ |
| 4C | four genotypes | 5 | $2\lambda_{\text{opt}}\tau_0$ | 101 | 1 | $\leq 4 \times 10^4$ | none |
| 4D | rare mutant | 2–16 | $\log 7$ | 101 | 20 | $\leq 2 \times 10^5$, tail $5 \times 10^3$ | none |
| 4E | evolution, independent | 5, 8, 12 | 3.5 | 101 | 2–6 | $5 \times 10^5$, tail $2.5 \times 10^5$ | $\sigma_{m,\phi} = 0.03$, $\sigma_{m,\sigma_f} = 0.02$, $\sigma_{m,\sigma_q} = 0.2$, $\tau_{\text{mut}} = 10$, $K = 50$, $\sigma_q \leq 10$ |
| 5 | evolution, independent | 5 | 3.5 | 101 | 8 per region | $5 \times 10^5$, tail $2.5 \times 10^5$ | as 4E |

#### C. Environmental noise depends on dilution and supply fluctuations

It is convenient to express the theoretical instability criterion in terms of the realized cycle growth factor. We therefore use

$$\sigma_\zeta^2 = \text{Var}_{\text{stat}}\left[\frac{\delta^d - \bar{\delta}}{\bar{\delta}}\right], \qquad \bar{\delta} = \mathbb{E}_{\text{stat}}[\delta^d], \tag{88}$$

where $\delta^d$ is the logarithmic biomass expansion required in cycle $d$, measured along the resident biomass trajectory. This definition is independent of whether the fluctuations originate from dilution, resource supply, or both, and it is the quantity that enters the local branching thresholds derived above.

Using Eq. (22), the unnormalized variance of the growth factor can be written as

$$\begin{aligned} \text{Var}_{\text{stat}}(\delta^d) = \text{Var}(\log D^d) + \text{Var}_{\text{stat}}\left(\log \frac{N_{\text{res}}^{d+1}}{N_{\text{res}}^d}\right) \\ + 2\,\text{Cov}_{\text{stat}}\left(\log D^d, \log \frac{N_{\text{res}}^{d+1}}{N_{\text{res}}^d}\right). \end{aligned} \tag{89}$$

Thus the imposed physical noise and the realized $\sigma_\zeta$ are not identical: biomass carry-over filters dilution and supply fluctuations before they enter the environmental-noise coordinate.

For a homogeneous resident that fully depletes the supplied resource, total biomass obeys

$$N^{d+1} = \frac{N^d + c_0^d}{D^d}, \tag{90}$$

$$\delta^d = \log\left(1 + \frac{c_0^d}{N^d}\right). \tag{91}$$

Combining this map with Eq. (22) gives the realized growth factor generated by the resident process. Equivalently,

$$\log \frac{N^{d+1}}{N^d} = \log\left(1 + \frac{c_0^d}{N^d}\right) - \log D^d. \tag{92}$$

Thus the current growth factor depends on the current supply and on the biomass carried over from previous dilution and supply draws.

This filtering can be computed explicitly in the low-noise regime. Let $N_\star = \bar{c_0}/(e^{\bar{\delta}} - 1)$ be the deterministic resident biomass and write

$$\eta^d = \log D^d - \bar{\delta}, \qquad \xi^d = \frac{c_0^d - \bar{c_0}}{\bar{c_0}}, \qquad n^d = \frac{N^d - N_\star}{N_\star}. \tag{93}$$

Linearizing Eq. (90) gives

$$n^{d+1} = e^{-\bar{\delta}} n^d + \left(1 - e^{-\bar{\delta}}\right) \xi^d - \eta^d, \qquad \delta^d - \bar{\delta} = \left(1 - e^{-\bar{\delta}}\right)\left(\xi^d - n^d\right). \tag{94}$$

If dilution and supply draws are temporally independent and cross-covariances are neglected, the stationary variance of the realized growth factor is

$$\mathrm{Var}_{\mathrm{stat}}(\delta^d) \simeq \frac{e^{\bar{\delta}} - 1}{e^{\bar{\delta}} + 1} \mathrm{Var}(\log D^d) + \frac{2(e^{\bar{\delta}} - 1)^2}{e^{\bar{\delta}}(e^{\bar{\delta}} + 1)} \frac{\mathrm{Var}(c_0^d)}{\bar{c_0}^2}. \tag{95}$$

Equivalently, for small fluctuations in $C^d = \log c_0^d$,

$$\mathrm{Var}_{\mathrm{stat}}(\delta^d) \simeq \frac{e^{\bar{\delta}} - 1}{e^{\bar{\delta}} + 1} \mathrm{Var}(\log D^d) + \frac{2(e^{\bar{\delta}} - 1)^2}{e^{\bar{\delta}}(e^{\bar{\delta}} + 1)} \mathrm{Var}(C^d). \tag{96}$$

The quantity to compare with the threshold in Eq. (47) is therefore

$$\sigma_\zeta^2 \simeq \frac{1}{\bar{\delta}^2}\left[\frac{e^{\bar{\delta}} - 1}{e^{\bar{\delta}} + 1} \mathrm{Var}(\log D^d) + \frac{2(e^{\bar{\delta}} - 1)^2}{e^{\bar{\delta}}(e^{\bar{\delta}} + 1)} \mathrm{Var}(C^d)\right]. \tag{97}$$

If supply is fixed, this reduces to

$$\sigma_\zeta^2 \simeq \frac{e^{\bar{\delta}} - 1}{e^{\bar{\delta}} + 1} \frac{\mathrm{Var}(\log D^d)}{\bar{\delta}^2}. \tag{98}$$

Thus constant supply, $\log c_0^d = \log \bar{c_0}$, removes direct supply noise but does not make the growth factor variance equal to $\mathrm{Var}(\log D^d)$. The imposed dilution variance is recovered only in the limit of large growth factor or when environmental noise is imposed directly. If both dilution and supply fluctuate, the limit of large growth factor gives instead $\sigma_\zeta^2 \simeq [\mathrm{Var}(\log D^d) + 2\mathrm{Var}(C^d)]/\bar{\delta}^2$.

The simulator controls the raw dilution noise rather than $\sigma_\zeta$. For fixed supply with gamma-distributed dilution, the imposed parameter is

$$\sigma_D = \frac{\mathrm{sd}(\log D^d)}{\bar{\delta}}, \qquad \log D^d \sim \Gamma\left(\sigma_D^{-2}, \bar{\delta}\, \sigma_D^2\right), \tag{99}$$

with shape $\sigma_D^{-2}$ and scale $\bar{\delta}\sigma_D^2$, so that $\mathbb{E}[\log D^d] = \bar{\delta}$. The imposed $\sigma_D$ and the realized $\sigma_\zeta$ differ because biomass carry-over filters the fluctuations (Eq. (89)), by a factor that depends on $\bar{\delta}$ and on the noise itself; Eq. (98) gives it only at small noise. Comparisons with the theory for fixed supply therefore use the resident map

$$\sigma_D \longmapsto \sigma_\zeta(\sigma_D) = \frac{\mathrm{sd}_{\mathrm{stat}}(\delta^d)}{\bar{\delta}}, \tag{100}$$

computed along the resident biomass trajectory, and a theoretical threshold $\sigma_{\zeta,\mathrm{crit}}$ corresponds to the imposed threshold $\sigma_D^{\mathrm{crit}}$ defined by

$$\sigma_\zeta(\sigma_D^{\mathrm{crit}}) = \sigma_{\zeta,\mathrm{crit}}. \tag{101}$$

For the reference dilution $D = 7$ the ratio $\sigma_\zeta/\sigma_D$ is close to 0.9 near the thresholds, so that comparing numerical thresholds in $\sigma_D$ with theoretical thresholds in $\sigma_\zeta$ would shift them systematically. In the Supplementary Material $\sigma_\zeta$ always denotes the measured environmental noise of Eq. (88), and the raw simulator input is written $\sigma_D$ when it is shown. In random-supply simulations the physical inputs $\sigma_D$ and the supply noise $\sigma_c$ are reported together with the measured $\sigma_\zeta$, which is the quantity compared with the theory.

#### D. Supply fluctuations enter through the cycle growth factor

For two resources with stochastic $D^d$ and stochastic $c_0^d$, the invasion criterion remains Eq. (27). The environmental input to this criterion is the environmental noise $\sigma_\zeta$. The low-noise expansion in Eq. (96) shows how fluctuations in $\log D^d$ and $\log c_0^d$ enter this environmental-noise coordinate.

We used fixed-strategy competition assays in the two-resource alternating environment. Both genotypes have the same mean allocation, $f = f_{\text{opt}}$. The resident is homogeneous, and the mutant is beta-distributed around the same mean with width $\sigma_f$. No mutation occurs during the assay. The imposed dilution noise is held fixed at $\sigma_D = 1.205$, for $\bar{\delta} = \log 7$. The supply is lognormal with coefficient of variation $\sigma_c$, the lag is glycolytic with $\lambda^\star = 1.1$, $\lambda_{\max} = 2.85$ and $\tau_0 = 1$, the mutant is represented by $N_{\text{pheno}} = 201$ phenotypes, and each condition has 50 stochastic histories. With fixed supply, the resident trajectory gives $\sigma_\zeta \simeq 1.09$, below the fixed-supply threshold $\sigma_{\zeta,\text{crit}} \simeq 1.19$ (Eq. (50)). Adding supply fluctuations with coefficient of variation $\sigma_c = 4$ increases the measured environmental noise to $\sigma_\zeta \simeq 1.22$.

The outcome is scored from the log-ratio $\ell^d$ of Eq. (86), after an initial transient. At the same imposed dilution noise, fixed supply eliminates the heterogeneous mutant, whereas stochastic supply makes the same mutant dominate over the assay horizon (Fig. 3).

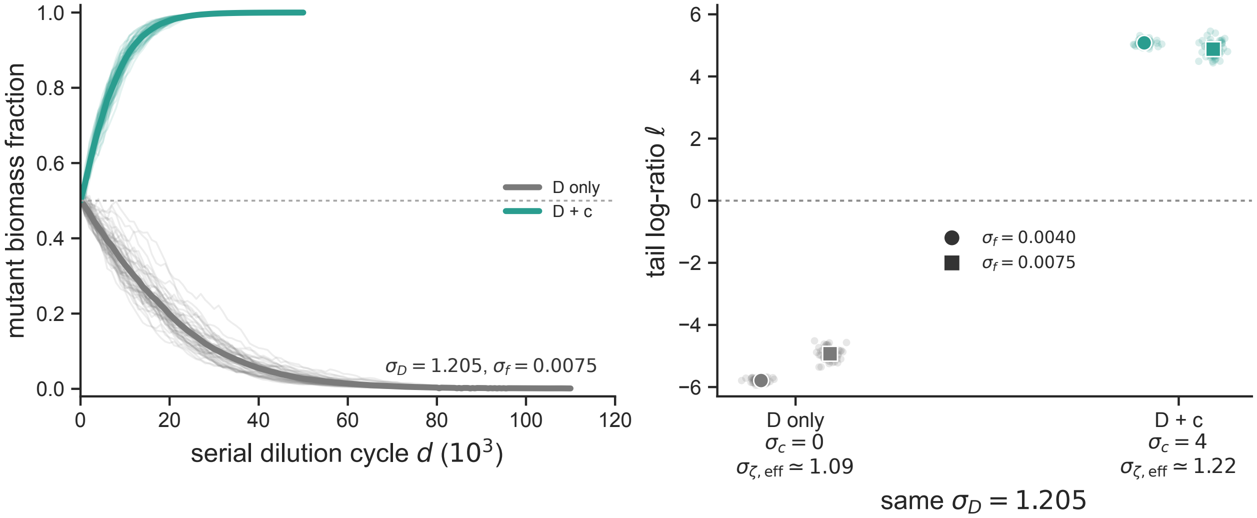


FIG. 3. **Random supply can make a below-threshold dilution environment favor phenotypic heterogeneity.** The imposed dilution noise is held fixed. Left, mutant biomass fraction through serial-dilution cycles for a heterogeneous mutant with $\sigma_f = 0.0075$. Thin curves are individual stochastic histories and thick curves are means across histories. With fixed supply, $\sigma_c = 0$, the measured environmental noise is $\sigma_\zeta \simeq 1.09$ and the mutant is eliminated. With stochastic supply, $\sigma_c = 4$, the measured environmental noise is $\sigma_\zeta \simeq 1.22$ and the mutant dominates over the assay horizon. Right, tail log-ratio $\ell$ for two levels of mutant variability. Translucent points are individual histories; symbols and bars are means and standard errors. Positive $\ell$ indicates heterogeneous-mutant advantage. The same dilution-noise background therefore selects opposite strategies depending on whether supply fluctuations move the resident growth-factor process across the threshold.

#### E. The four regions of the map hold for the gluconeogenic lag and shift for non-symmetric transitions

For the glycolytic lag the four regions of the map depend on a single number, $\lambda_{\text{opt}}\tau_0$. For the gluconeogenic lag the lag cost depends on the target of the switch, so the boundaries are expected to depend on the transition matrix (Sec. IV E): for symmetric transitions the map keeps its structure with $\lambda_{\text{opt}}\tau_0$ replaced by $x = \lambda_{\text{opt}}\tau_0/r^{\text{opt}}$, while for non-symmetric transitions the boundary of all-in hedging is predicted to move with the predictability of the switches (Eq. (74)). We test both predictions.

We run the four-genotype competitions of main Fig. 4C (Table I) for the gluconeogenic lag: a homogeneous genotype and genotypes with variability in $f$, in $q$ and in both, sharing the optimal mean allocation of the homogeneous resident, without mutations. Transitions among the $M = 5$ resources are circulant, the switch from $b$ going to $b + k$ with probability $w_k$; we consider the symmetric matrix and three matrices in which the probability concentrates progressively on one target. For each matrix, points of the plane $(x, \sigma_\zeta)$ are sampled at random and realized by choosing $\tau_0$, $\bar{\delta}$ and the imposed dilution noise. The levels of variability, $\sigma_f = 0.02$ and $\sigma_q = 0.05$, are small, as the local theory requires; this pair was chosen among a small grid as the one that best reproduces the symmetric map, and is used for all matrices.

For symmetric transitions the winner is the one predicted by the local theory at nearly all decided points (98%); the exceptions lie near the boundary of the all-in term, where the genotype with variability in both $f$ and $q$ wins instead of the one with variability in $f$ only, a finite-amplitude effect as for the glycolytic lag (Fig. 4). For non-symmetric transitions the four regions persist: the boundary of the all-in term moves to smaller $x$ as the probability concentrates on one target, as predicted to within 2%, while the boundary of the growth term, $\sigma_\zeta^2 x = 2$, does not move. A more concentrated matrix is not used, because the lag of its dominant target is clipped at zero for the resident, where the local expansion does not apply.

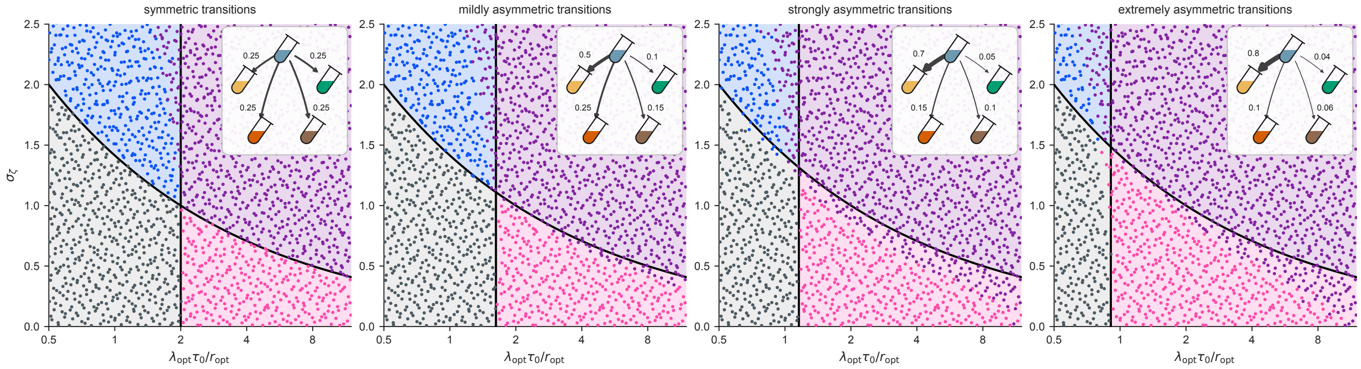


FIG. 4. **For the gluconeogenic lag the local regions hold, and the boundary of the all-in term moves with the predictability of the transitions as predicted.** Winner of the competition between a homogeneous genotype and genotypes with variability in $f$, in $q$ and in both, in the plane $(\lambda_{\rm opt}\tau_0/r^{\rm opt}, \sigma_\zeta)$, for $M = 5$ and circulant transitions whose probabilities are shown in the insets (arrow width proportional to the probability), from symmetric to extremely asymmetric; colors as in main Fig. 4C. Lines and shaded regions are the boundaries of the local theory, $\sigma_\zeta^2 \lambda_{\rm opt}\tau_0/r^{\rm opt} = 2$ and Eq. (74). 2048 points per panel, $\sigma_f = 0.02$, $\sigma_q = 0.05$, $N_{\rm pheno} = 101$.

#### F. The evolved variability does not depend on the supply of mutations

The analytical results of the main text follow from the invasion of a resident at its stationary state by a rare mutant (Sec. II). They describe the successional regime, in which each mutant either invades or is excluded before the next one arrives. In the evolutionary simulations, instead, mutants are introduced every $\tau_{\rm mut} = 10$ cycles on average (Table I), whereas the sweep of a mutant with invasion fitness $s \simeq 2 \times 10^{-3}$, measured from the first sweeps of the runs, takes about $\ln(10^7)/s \simeq 8 \times 10^3$ cycles to go from its initial abundance to dominance. Many mutants therefore grow at the same time, and several genotypes can be close to dominance simultaneously (Fig. 5A). In this regime of interference between concurrent sweeps the evolved state is not guaranteed to coincide with the one selected by a sequence of independent invasions. We test whether it depends on the supply of mutations by varying the mutation interval $\tau_{\rm mut}$ with all other settings fixed, and with protocols that limit the number of mutants that are undecided at the same time.

All runs use two resources with $\sigma_D = 1.5$, above the threshold $\sigma_{D,\rm crit} \simeq 1.19$, fixed supply, the one-axis mutation kernel of main Fig. 3 and the lineage cap $K = 50$ (Table II). Three values of the mutation interval are compared with three replicates each, starting from the homogeneous optimum with $\sigma_f = 0.03$: $\tau_{\rm mut} = 10$, strong interference, with tens of lineages above $10^{-3}$ at the same time; $\tau_{\rm mut} = 10^3$, weak interference, where sweeps overlap only partially; and $\tau_{\rm mut} = 4 \times 10^4$, where mutations are rare enough that the first sweeps occur one at a time. Two controls hold the number of undecided mutants fixed instead of the interval. In the successional protocol a new mutant is introduced only when the biomass outside the dominant lineage is below $10^{-8}$ of the total, that is, when the previous mutant has been excluded or has fixed and the former resident has disappeared; to avoid blocking the run when a mutant is nearly neutral, a new mutant is introduced anyway after $2 \times 10^5$ cycles without a decision. In the second control up to $n_{\rm conc} = 3$ or 10 undecided mutants are allowed at the same time, with the same criterion. We also repeat the strong-interference run with the lineage cap $K = 200$. Finally, one run per regime starts from each of four initial conditions, $(f, \sigma_f) \in \{0.5, 0.6\} \times \{0.03, 0.004\}$.

Mutation supply changes the dynamics but not the state reached. In the strong-interference runs the dominant genotype is replaced every $10^4$ cycles or so while tens of lineages rise simultaneously, and the variability falls below 0.012 after about $10^5$ cycles; with rarer mutations the takeovers become sequential and the same level is reached after about $4 \times 10^5$ cycles at $\tau_{\rm mut} = 10^3$ and $2 \times 10^6$ at $\tau_{\rm mut} = 4 \times 10^4$ (Fig. 5A–D). In the successional protocol it takes about $5 \times 10^6$ cycles. At long times all protocols reach the same variability, $\bar{\sigma}_f^B = 0.008$–$0.010$, with the mean allocation within 0.004 of $f_{\rm opt} = 0.5207$ (Table II), and the runs started from different $(f, \sigma_f)$ converge to the same state in the three regimes (Fig. 5E). Larger lineage caps do not change the endpoint either.

Since the final state does not depend on the supply of mutations, whereas the time to reach it grows by two orders of magnitude when mutations are rare, in the simulations of the main text we use $\tau_{\rm mut} = 10$, which gives evolutionary convergence in $10^5$–$10^6$ cycles instead of $10^7$.

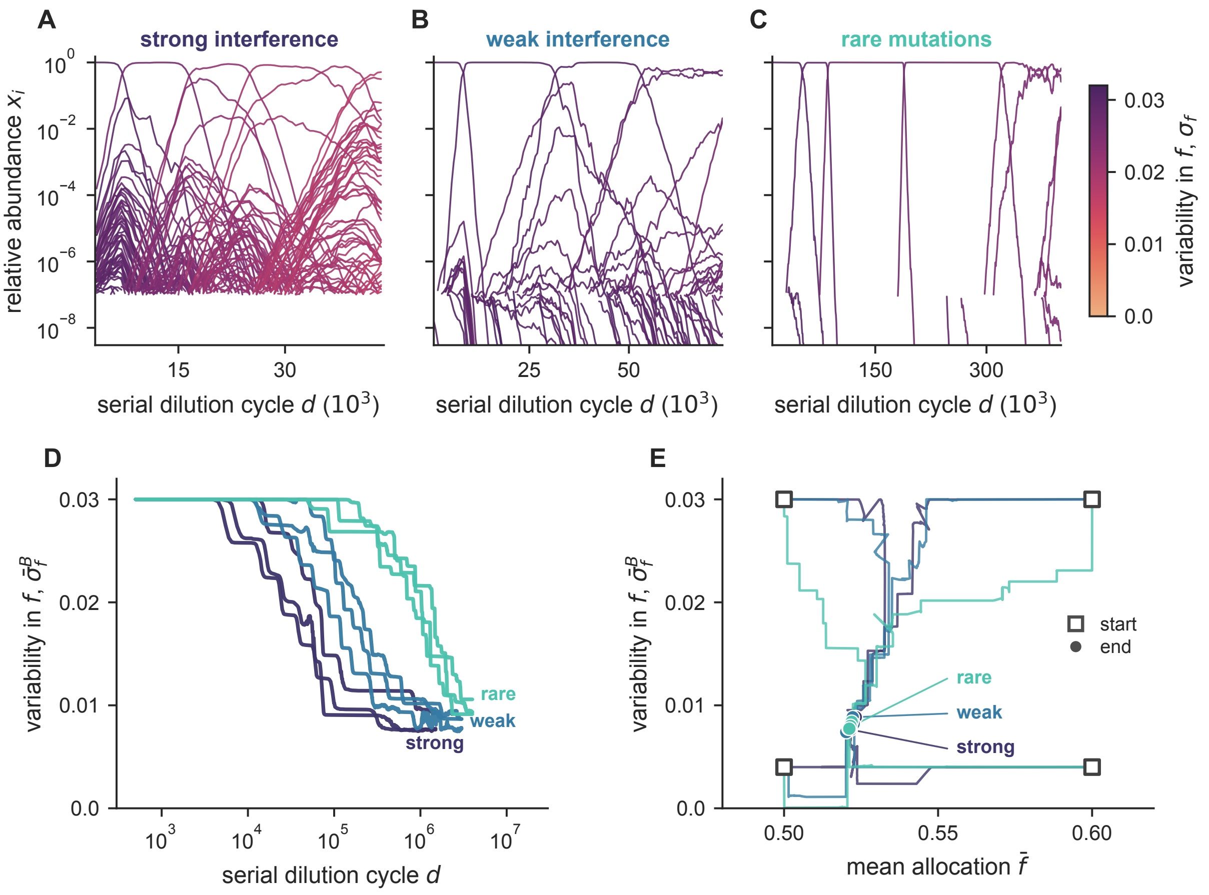


FIG. 5. **The evolved variability and mean allocation do not depend on the supply of mutations, whereas the time needed to reach them does.** (A-C) Relative abundance of every lineage in one run per regime, in a window containing the takeover of four successive genotypes; the window differs among panels: strong interference ($\tau_{\rm mut} = 10$), weak interference ($\tau_{\rm mut} = 10^3$) and rare mutations ($\tau_{\rm mut} = 4 \times 10^4$). Colour is the variability $\sigma_f$ of each lineage. (D) Biomass-weighted variability $\bar{\sigma}_f^B$ versus serial dilution cycle for the three replicates of each regime. (E) Trajectories in the plane of the mean allocation and the variability from four initial conditions, with one run per regime; squares mark the initial conditions and circles the final states. All runs have $\sigma_D = 1.5$ and mutations in the mean allocation or in $\sigma_f$ (one-axis kernel).

TABLE II. **Evolved state for different supplies of mutations.** Mean and standard deviation over replicates of the biomass-weighted variability $\bar{\sigma}_f^B$ and mean allocation $\bar{f}$ over the last 20% of each run ($\sigma_D = 1.5$, one-axis mutations, start at $f_{\rm opt}$ and $\sigma_f = 0.03$, $M = 2$, $N_{\rm pheno} = 201$, $K = 50$ unless stated). Kept is the fraction of introduced mutants that survive for more than 100 cycles.

| protocol | cycles | runs | $\bar{\sigma}_f^B$ | $\bar{f}$ | mutants | kept |
|---|---|---|---|---|---|---|
| $\tau_{\rm mut} = 10$ | $1.5 \times 10^6$ | 3 | $0.0082 \pm 0.0010$ | $0.5220 \pm 0.0022$ | $1.5 \times 10^5$ | 3% |
| $\tau_{\rm mut} = 10$, $K = 200$ | $1.5 \times 10^6$ | 1 | 0.0090 | 0.5234 | $1.5 \times 10^5$ | 18% |
| $\tau_{\rm mut} = 10^3$ | $3 \times 10^6$ | 3 | $0.0084 \pm 0.0007$ | $0.5217 \pm 0.0007$ | $3.0 \times 10^3$ | 72% |
| $\tau_{\rm mut} = 4 \times 10^4$ | $4 \times 10^6$ | 3 | $0.0098 \pm 0.0007$ | $0.5241 \pm 0.0012$ | 100 | 100% |
| $n_{\rm conc} = 3$ | $2 \times 10^7$ | 3 | $0.0082 \pm 0.0006$ | $0.5220 \pm 0.0007$ | 190 | 100% |
| $n_{\rm conc} = 10$ | $6 \times 10^6$ | 3 | $0.0083 \pm 0.0004$ | $0.5218 \pm 0.0012$ | 286 | 100% |
| successional | $2 \times 10^7$ | 3 | $0.0087 \pm 0.0007$ | $0.5223 \pm 0.0009$ | 132 | 100% |

[1] D. W. Erickson, S. J. Schink, V. Patsalo, J. R. Williamson, U. Gerland, and T. Hwa, Nature **551**, 119 (2017).
[2] M. Basan, T. Honda, D. Christodoulou, M. Hörl, Y.-F. Chang, E. Leoncini, A. Mukherjee, H. Okano, B. R. Taylor, J. M. Silverman, *et al.*, Nature **584**, 470 (2020).
[3] O. Kotte, B. Volkmer, J. L. Radzikowski, and M. Heinemann, Molecular systems biology **10**, 736 (2014).
[4] F. Schreiber, S. Littmann, G. Lavik, S. Escrig, A. Meibom, M. M. Kuypers, and M. Ackermann, Nature microbiology **1**, 16055 (2016).
[5] R. M. Y. Omer, O. Mazzarisi, M. Dal Bello, and J. Grilli, Evolutionary learning of microbial populations in partially predictable environments (2025), arXiv:2508.00527 [q-bio.PE].
[6] A. M. New, B. Cerulus, S. K. Govers, G. Perez-Samper, B. Zhu, S. Boogmans, J. B. Xavier, and K. J. Verstrepen, PLoS biology **12**, e1001764 (2014).
